# A Reaction-Driven Condensate-to-Vesicle Transition Selects, Activates, and Spatially Organizes RNA

**DOI:** 10.64898/2026.06.07.730732

**Authors:** Hong-Guen Lee, Alessandro Fracassi, Alexander Harjung, Taeyang An, Neal K. Devaraj

## Abstract

Living systems depend on the selective concentration of informational polymers within membrane-bound compartments. Liquid-liquid phase-separated condensates efficiently concentrate biomolecules, whereas membrane-bound vesicles provide persistent compartment boundaries. However, direct chemical mechanisms that couple these organizational states remain largely unexplored. Here we report a reaction-driven pathway linking condensates to RNA-enriched lipid vesicles. Electrostatic interactions between cationic thioesters and RNA drive phase separation into reactive condensates. Reaction with cysteine generates membrane lipids and transforms the condensates into unilamellar bilayer vesicles with near-uniform size distribution. The vesicles encapsulate >90% of the RNA from the initial solution and concentrate it by more than two orders of magnitude to >100 µM. Encapsulation is strongly dependent on RNA length, with short oligonucleotides excluded while longer RNAs are selectively retained. Under spatial gradients of the chemical trigger, sharply defined vesicle populations with distinct RNA compositions emerge. Reaction-driven compartmentalization raises local ribozyme and substrate concentrations above the threshold required for catalytic activity, enabling function from otherwise inactive dilute solutions. These findings establish a mechanism by which chemical reactions generate selective, functional, and spatially organized RNA-enriched membrane-bound compartments from heterogeneous molecular mixtures.

## Introduction

Living systems rely on well-defined persistent boundaries that regulate and localize chemical reactions and maintain high concentrations of proteins and genetic polymers.^1–7^ However it remains unclear how hierarchical membrane-bound compartments enriched in biomolecules can spontaneously arise from mixtures of simple organic molecules, such as lipids, amino acids, and nucleic acids, especially under dilute conditions where intermolecular interactions are weak.^8–11^ Numerous studies have explored engineering biological compartmentalization.^12–17^ Lipid membranes define the boundaries of cells, and membrane-bound vesicles have frequently been studied because they provide well-defined yet deformable boundaries that act as semi-permeable barriers that isolate and sustain chemical reactions.^18–20^ However, vesicles are typically formed from preexisting amphiphiles by hydrating lipid films and encapsulate biomolecules such as nucleic acids inefficiently without external manipulation or specialized instrumentation.^21–25^ Alternatively, liquid-liquid phase-separated condensates or coacervates represent another major compartmentalization paradigm commonly used by biology.^26–31^ Such condensates spontaneously form, often through multivalent interactions, and can concentrate biopolymers like RNA and proteins, even from dilute starting solutions.^32– 35^ However, unlike lipid bilayers, liquid-liquid condensates lack robust barrier properties and provide limited control over molecular exchange, making it difficult to retain concentrated components, sustain chemical gradients, and support durable compartmentalized function.^32,34,36,37^ As lipid bilayer and condensate compartmentalization strategies typically require assembly from different classes of molecular building blocks, they have largely been explored independently.^12,27,38^

The benefits and drawbacks of different compartmentalization approaches are particularly important when encapsulating genetic polymers such as RNA. Elevated RNA concentrations stabilize molecular information against stochastic loss and promote the intermolecular interactions required for RNA catalysis and cooperative folding.^37–39^ At the same time, the compartment must have robust boundaries so that nucleic acids do not exchange.^3,42,43^ Finally, the compartments themselves should be of uniform size to enable reproducible internal chemistry across the population, which would be a prerequisite for effective selection and evolution.^2,3,42^ Identifying a spontaneous compartmentalization mechanism that meets these requirements has been an unmet challenge.

Here we report a chemically gated pathway that starts from dilute solutions of thioesters, RNA, and amino acids and results in the spontaneous formation of nearly uniformly sized lipid compartments that are highly enriched in RNA. Electrostatic interactions between RNA and cationic thioesters drive liquid–liquid phase separation forming reactive condensates. The addition of the amino acid cysteine causes a spontaneous chemical transformation within these condensates, producing diacyl membrane lipid products. Lipid synthesis triggers the transformation of the condensates into unilamellar bilayer vesicles with a narrow size distribution centered between 1–2 µm depending on the length of the RNA. RNA size–dependent selection occurs during encapsulation, as very short RNAs (10 nt) are not captured, even in the presence of longer RNAs. Size-selective encapsulation can be further tuned by local cysteine concentration and spatial patterning occurs in the presence of a cysteine gradient, giving rise to sharply divided vesicle populations with distinct RNA compositions. The condensate-to-vesicle transition can capture over 90% of the RNA present in the initial solution and concentrate it by more than two orders of magnitude within the vesicles, yielding internal RNA concentrations exceeding 100 µM. The final vesicle-encapsulated RNA is protected from external stresses like high salt concentrations and degradative enzymes. RNA ribozymes and substrates concentrated during protocell formation show a dramatic increase in reactivity. Together, these results establish a physicochemical route by which chemical reactions couple compartment formation, molecular selection, and catalytic activation, transforming simple molecular mixtures into functional, RNA-enriched, membrane-bound compartments.

## Results

### Formation of RNA-C_12_CT Condensates and Vesicle Generation

We previously demonstrated spontaneous diacylation between positively charged choline thioester (CT) detergents and cysteine, resulting in the formation of membrane lipids.^44,45^ Micelles formed from positively charged detergents have been shown to condense with anionic polymers forming liquid-liquid phase-separated coacervates, suggesting that CTs could form condensates with anionic nucleic acid polymers like RNA.^46–50^ Electrostatic condensation between RNA and CTs was initially tested using 80 nt tRNA (1.3 µM final concentration) as a model oligoRNA. The addition of Quantifluor, an RNA-sensitive intercalating dye, enabled fluorescence microscopy visualization and a linear fluorescence readout of RNA concentration. C_12_CT (4 mM final concentration), a CT derivative bearing a dodecyl chain, was introduced along with 0.05 mol% DiI as a membrane-staining dye. Upon mixing in a microscopy chamber (Figure 1a), we observed uniform layers of phase-separated material on the glass surface in which both the Quantifluor and DiI signals colocalized, suggesting formation of a homogeneous condensate between RNA and the C_12_CT detergent (Figure 1b). FRAP (fluorescence recovery after photobleaching) measurements supported a liquid-liquid phase-separated condensate, as fluorescence recovery slowly took place after photobleaching of the RNA-binding dye (Figure S1). The phase diagram of 80 nt RNA and C_12_CT shows that condensate formation occurs over a defined range of RNA and detergent concentrations (Figure S2).

**Figure 1.**
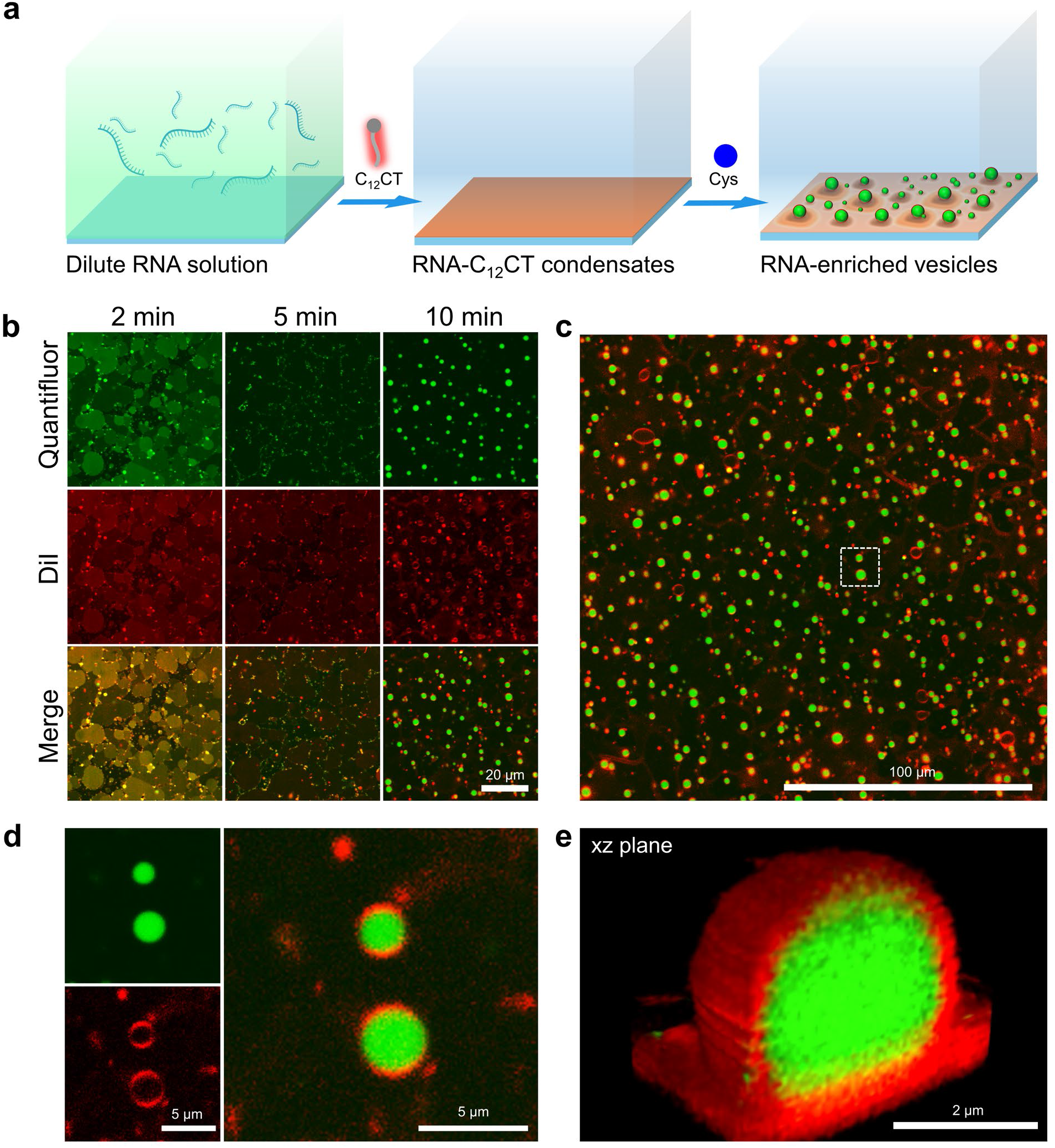
Chemically gated formation of RNA-enriched vesicles. a) Schematic of the formation of RNA-enriched vesicles from dilute RNA solutions. b) Time-lapse confocal fluorescence images of RNA-C_12_CT condensates stained with Quantifluor and DiI, acquired 2, 5, and 10 min after cysteine treatment. c) Wide-field image of RNA-enriched vesicles obtained 10 min after cysteine treatment. d) Magnified view of RNA-enriched vesicles. e) 3D reconstructions of a representative vesicle shown in the xz plane. All experiments were performed using 80 nt RNA.

We next investigated whether C_12_CTs in the RNA-C_12_CT condensates would undergo cysteine-driven diacylation and how this transformation affects the associated RNA. Treatment with cysteine (10 mM final concentration) caused the RNA-C_12_CT layers to contract into smaller domains and droplets while maintaining colocalization of Quantifluor and DiI signals (Figure 1b, Supplementary video 1). However, within 10 min, the DiI signal outlined circular membrane boundaries, while Quantifluor was confined to the enclosed interior. (Figure 1b, Supplementary video 1). These features are consistent with the formation of unilamellar vesicles composed of diacylated cysteine lipids, similar to previous studies.^44,45^ LCMS measurements confirmed diacylation of cysteine, with concomitant loss of C_12_CT (Figure S3). However, unlike prior studies, the vesicles observed on the glass surface appeared nearly uniform in size, with an average diameter of 1.5 ± 0.29 µm (Figure 1c, S9). 3D reconstructions further supported vesicle unilamellarity and revealed spherical vesicles containing concentrated RNA, as indicated by strong Quantifluor signal enclosed by DiI-stained membranes (Figure 1d–e). Based on the Quantifluor fluorescence intensity, a 102 ± 6-fold increase in intravesicular RNA signal, relative to the initial RNA solution, was observed after 10 min of cysteine treatment (Figure S4), corresponding to an RNA concentration of ~130 µM (Figure S5). An encapsulation-efficiency assay measuring total encapsulated RNA by urea–PAGE indicated that approximately 96 ± 8 % of the total RNA in the initial solution was encapsulated within the same time frame (Figure S6). RNA diffusion in the final vesicles was too fast to be measured by FRAP, supporting a highly solvated state in the interior of the vesicle that is in the same phase as the exterior aqueous solution (Figure S1). As expected, vesicles formed by hydration of preformed diacylated cysteine lipid films in the presence of the same concentration of RNA showed only background Quantifluor fluorescence due to extremely low encapsulation efficiency. The observed RNA enrichment requires cysteine-triggered conversion of RNA-C_12_CT condensates into vesicles (Figure S7), demonstrating that in situ lipid synthesis is essential.

### RNA-Enriched Vesicles Emerge via Condensate Intermediates

RNA-C_12_CT condensates change their physical properties as their intermolecular interactions evolve during cysteine diacylation. Loss of the positively charged C_12_CT thioester during diacylation would be expected to destabilize the condensate. Indeed, addition of 0.1 equivalents of cysteine led to an approximate 2700-fold increase in the diffusivity of Quantifluor-labeled RNA within the intermediate droplet-like domains, as determined by FRAP (Figure S1). Under such cysteine-deficient conditions (0.1–0.4 equivalents relative to C_12_CT), we also observed RNA-lipid condensates encapsulated within vesicles, suggesting that membrane formation initiates at the condensate surface, trapping RNA that was previously in the condensate phase. (Figure 2a,b). Additional experiments employing reduced amounts of C_12_CT (1–50% of the standard 4 mM condition) also highlighted the importance of lipid synthesis in generating RNA-enriched vesicles. When using 20-50% C_12_CT, the resulting vesicles initially encapsulated enriched RNA, but within 20 min of cysteine treatment most vesicles lost their RNA, likely due to disruption in membrane integrity. At 10% C_12_CT or lower, the RNA-C_12_CT condensates did not form vesicles (Figure S8). Super-resolution imaging further revealed that RNA-encapsulating vesicles originate directly from the RNA-thioester condensate droplets observed shortly after the addition of cysteine (Figure 2c). Together, these findings suggest that intermediate RNA-lipid phases form during the transition from condensates to vesicles, and that stabilizing the intermediate phase is essential for generating RNA-enriched vesicles. Based on these observations, we suggest a putative mechanism in which RNA and CT initially forms a dense liquid condensate characterized by slow diffusion. Cysteine-triggered diacylation consumes the positively charged C_12_CT detergent, weakening RNA electrostatic binding and giving rise to a more fluid intermediate condensate in which molecular diffusion is markedly increased. The diacylated amphiphilic cysteine likely assembles at the condensate–water interface,^51–53^ destabilizing the initially homogeneous condensate film and driving its breakup into discrete droplets. The resulting droplets exhibit a narrow size distribution, consistent with reaction-driven changes in interfacial properties that limit uncontrolled growth and coalescence.^54–57^ As thioester is continually transformed into diacylated cysteine lipids, the increased hydrophobicity at the condensate interface promotes continued segregation of lipid products, driving bilayer membrane formation, disruption of the condensate phase, and ultimately generating vesicles that capture the RNA that was originally sequestered in the condensate phase (Figure 2d).

**Figure 2.**
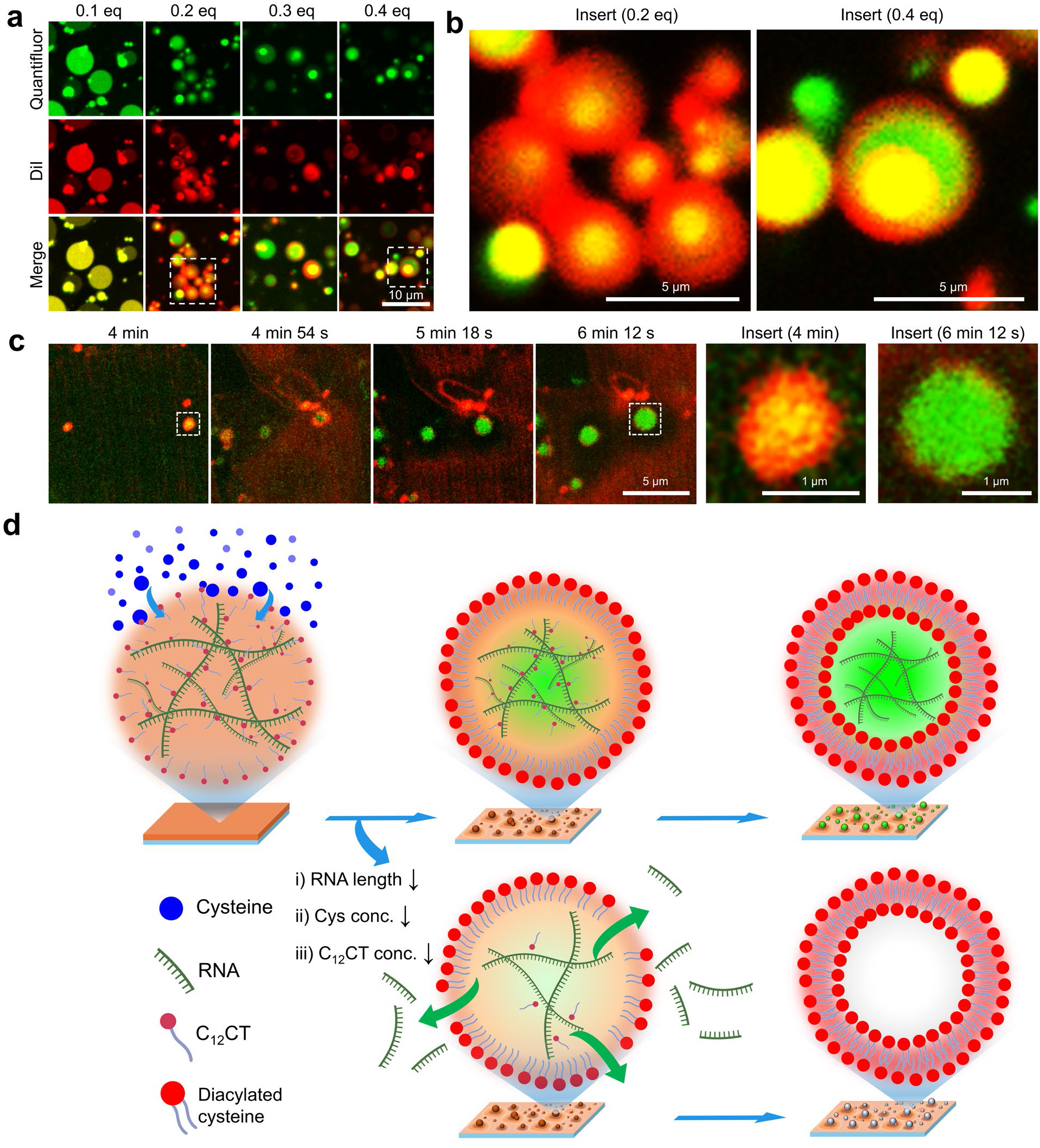
Formation of RNA-enriched vesicles via condensate intermediates. a) RNA-thioester condensates formed by mixing RNA-C_12_CT complexes with 0.1-0.4 equivalents of cysteine (relative to C_12_CT) and incubating for 12 h at 37 °C. b) Magnified views of the 0.2 and 0.4 equivalent conditions shown in panel a. c) Time-lapse super-resolution confocal images showing the emergence of highly fluid condensate intermediates (4 min) and their transition into RNA-enriched vesicles (4 min 54 s - 6 min 12 s). d) Schematic illustrating the transition from RNA-C_12_CT condensates to RNA-enriched vesicles. Cysteine-triggered diacylation converts the positively charged detergent to a negatively charged lipid, weakening RNA electrostatic binding and giving rise to highly fluid condensate droplet intermediates surrounded by diacylated cysteine. Increased hydrophobicity at the condensate surface promotes segregation of lipid precursors to the interface, driving membrane formation and ultimately generating vesicles that encapsulate concentrated RNA in an aqueous interior. Under conditions of insufficient RNA length, suboptimal cysteine levels, or low C_12_CT concentrations, RNA-C_12_CT condensates fail to remain stable during membrane lipid synthesis. All experiments were performed using 80 nt RNA.

### RNA is Size-Selectively Encapsulated in Vesicles

To further investigate how RNA length influences RNA-enriched vesicle formation, we repeated the experiments using RNAs of different lengths (10, 22, 447, and 976 nt in place of 80 nt RNA). Condensates containing RNAs from 22 nt to 976 nt generated RNA-enriched vesicles upon cysteine treatment, similar to those obtained with 80 nt RNA (Figure 3a). Vesicle size measured after 10 min of cysteine treatment showed a logarithmic dependence on RNA length, yielding average diameters of 1.1 μm, 1.5 μm, 1.6 μm, and 2.0 μm for 22 nt, 80 nt, 447 nt, and 976 nt RNAs, respectively (Figure S9). RNA encapsulation efficiency and retention were strongly dependent on RNA length. The 976 nt RNA showed lower enrichment (67 ± 6.7 fold) than the 80 nt RNA (102 ± 6.2 fold) but higher retention after 1 h (76% versus 67%), consistent with reduced membrane permeation of longer RNA polymers (Figure S10). In contrast, 22 nt RNA showed only 10 ± 1.1-fold enrichment and 15% retention under the same conditions, reflecting weaker complexation with CT and more rapid escape across the lipid membrane as it forms (Figure S11). Notably, these RNA length– dependent changes in vesicle size and encapsulation efficiency occurred despite a constant total amount of RNA in the initial solution. This behavior suggests that RNA polymer length influences the structure or dynamics of condensate intermediates, thereby biasing subsequent membrane assembly toward vesicles of different sizes. Longer RNA polymers may stabilize condensates during amphiphile synthesis, leading to larger precursor droplets and ultimately larger vesicles with reduced RNA leakage.

**Figure 3.**
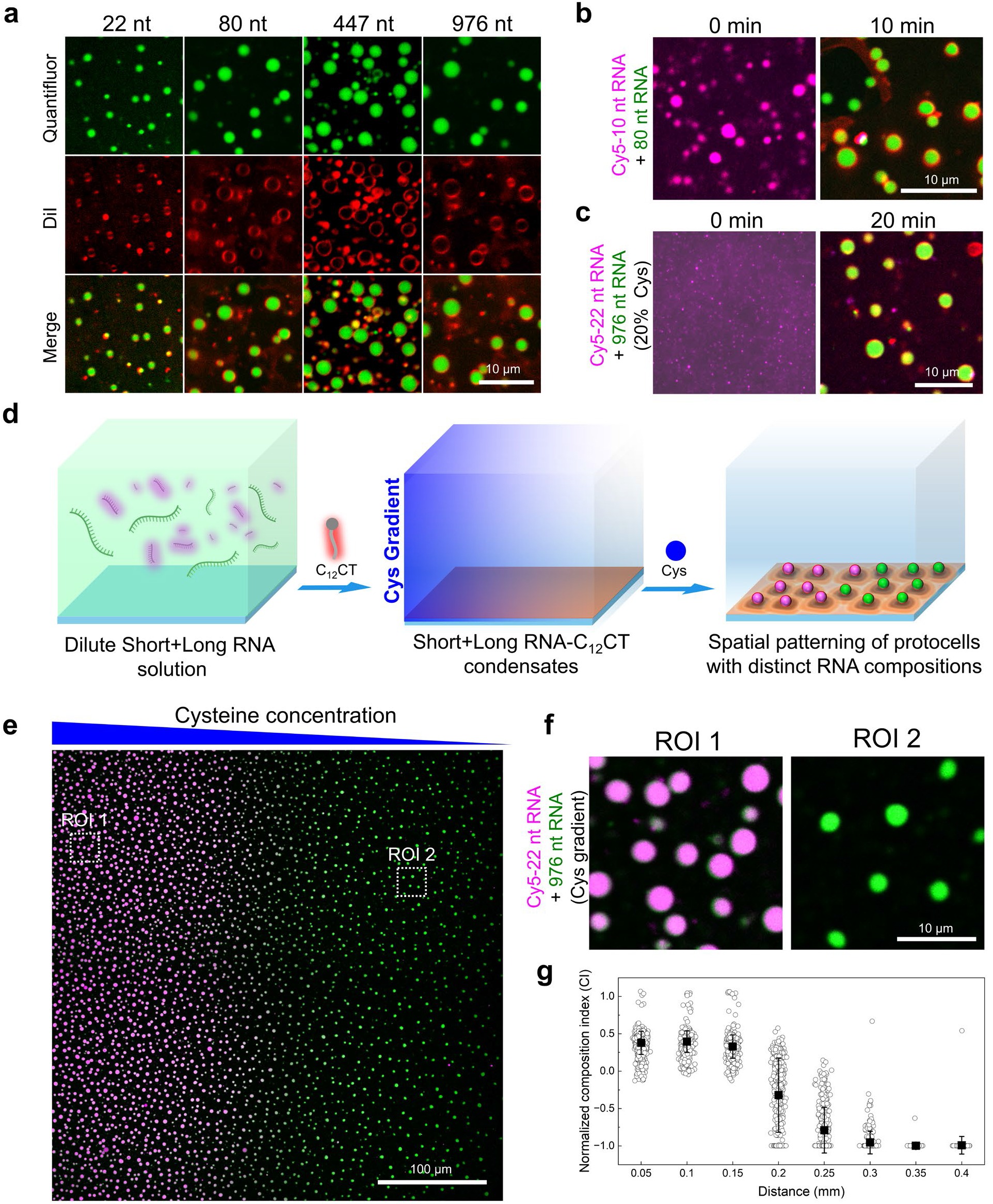
RNA Length-Dependent Selection Drives Spatial Differentiation of Vesicle Populations. a) Confocal fluorescence microscopy images of vesicles encapsulating 22, 80, 447, and 976 nt RNAs, acquired 10 min after cysteine addition to the corresponding RNA-C_12_CT condensates. b) Confocal fluorescence microscopy images of RNA-C_12_CT condensates containing a 1:1 (w/w) mixture of Cy5-10 nt RNA and 80 nt RNA and of RNA-enriched vesicles derived from the corresponding condensates 10 min after cysteine addition. c) Confocal fluorescence microscopy images of RNA-C_12_CT condensates containing a 1:1 (w/w) mixture of Cy5-22 nt RNA and 976 nt RNA and of RNA-enriched vesicles formed from the corresponding condensates 20 min after addition of 20% cysteine relative to the original condition. In b and c, images are shown as merges of Cy5-labeled RNA (magenta), Quantifluor (green), and DiI (red). Individual channels are shown in Figure S15 and S17. d) Schematic showing the cysteine gradient-driven patterning of vesicles with distinct RNA compositions from a dilute solution containing short and long RNA species. e) Confocal fluorescence microscopy image of spatially patterned vesicles with distinct RNA compositions, acquired 20 min after cysteine addition to RNA-C_12_CT condensates consisting of a 1:1 (w/w) mixture of Cy5-22 nt RNA and 976 nt RNA. Cysteine solution was added to the left edge of the chamber to establish a left-to-right cysteine gradient. Magenta (Cy5-22 nt RNA) and green (Quantifluor) channels are merged. The DiI channel is omitted for clarity. f) Magnified views of ROI 1 and ROI 2 in e, showing vesicles near the left and right edges of the image, respectively. Individual channels, including DiI, are shown in Figure S18. g) Normalized composition index (CI) showing the change in the relative intensities of Cy5-22 nt RNA and Quantifluor fluorescence in panel e. CI values of 1 and −1 indicate the presence of only Cy5-22 nt RNA signal or only Quantifluor signal, respectively. Distance on the x-axis indicates distance from the left edge of the field of view shown in panel e.

In contrast, shorter RNAs fail to remain associated with vesicles during the condensate-to-vesicle transition. After addition of cysteine to 10 nt RNA-C_12_CT condensates, RNA was observed to escape at approximately 5 min and was completely lost by 10 min (Figure S12). These vesicles were larger and more heterogeneous in size and shape (Figure S12), and often multilamellar, resembling structures observed during in situ lipid synthesis in the absence of RNA.^44,45^ LCMS analysis showed similar diacylation kinetics for 10 nt and 80 nt RNA-C_12_CT condensates (Figure S3 and S13), indicating that RNA retention is governed by RNA length rather than lipid reaction kinetics, likely because longer RNAs help sustain condensate stability even as the thioester is depleted during the reaction. Similarly, 15 nt RNA-C_12_CT condensates showed only transient enrichment into submicron droplets (Figure S14). In mixtures of Cy5-10 nt RNA and 80 nt RNA, the Cy5 fluorescence signal rapidly dissipated and was lost from the droplets and vesicles upon cysteine treatment, whereas the Quantifluor signal still accumulated within vesicles, suggesting selective enrichment of the longer 80 nt RNA (Figure 3b and S15). RNase A treatment (10^-5^ mg mL^-1^) of the 10 nt and 80 nt mixed RNA-enriched vesicles followed by urea-PAGE verified selective length-dependent enrichment of the 80 nt RNA (Figure S16). We examined whether selective enrichment of the longer RNA could also occur in other RNA mixtures. When RNA-C_12_CT condensates containing Cy5-22 nt RNA and 976 nt RNA were treated with 10 mM cysteine, they gave rise to vesicles containing both 22 nt and 976 nt RNAs, as evidenced by colocalized Quantifluor and Cy5 signals within the vesicles (Figure S17). However, when the cysteine concentration was reduced to 2 mM, vesicle formation slowed, and after 20 min the resulting vesicles were selectively enriched in the 976 nt RNA, as indicated by Quantifluor fluorescence within vesicle interiors and the absence of Cy5 signal (Figure 3c and Figure S17). These results show that selective encapsulation of the longer RNA in a heterogeneous RNA mixture can be promoted not only by RNA size differences but also by changing the cysteine concentration, which modulates vesicle-formation kinetics.

### Cysteine Gradients Drive Spatial Patterning of Vesicles with Distinct RNA Compositions

The role of cysteine concentration in size-selective RNA encapsulation led us to investigate whether a cysteine gradient could direct the spatial patterning of vesicles with distinct RNA compositions, analogous to the way morphogen gradients spatially organize homogeneous cell populations into distinct cell groups (Figure 3d).^58–61^ To examine the spatial patterning of vesicles with distinct RNA compositions, we repeated the RNA-enriched vesicle formation experiment using a mixture of Cy5-22 nt RNA and 976 nt RNA, with cysteine (10 mM final concentration) added to the left edge of the chamber to establish a left-to-right concentration gradient. At 20 min after cysteine addition, the resulting vesicles displayed spatially distinct RNA compositions along the gradient, with vesicles on the left encapsulating both 22 nt and 976 nt RNAs (Figure 3f and S18, ROI 1), whereas those on the right encapsulated only 976 nt RNA (Figure 3f and S18, ROI 2). Interestingly, we observe a sharp boundary reminiscent of French flag patterning (Figure 3e).^62–64^

The left-to-right shift in composition index (CI) from 0.3 to −1, accompanied by a narrow standard deviation, supports a collective and coherent change in RNA composition across thousands of vesicles within a submillimeter-scale region, demonstrating that the cysteine gradient drives spatial patterning of vesicles with distinct RNA compositions (Figure 3g). Time-lapse imaging revealed the simultaneous emergence at 5 min of vesicles containing both Cy5-22 nt and 976 nt RNAs on the left side of the chamber, together with the loss of Cy5-22 nt RNA signal and the subsequent appearance at 7.5 min of vesicles enriched only in 976 nt RNA on the right side, suggesting that spatial patterning arises during the condensate-to-vesicle transition due to differences in vesicle-formation kinetics (Figure S19). Similar spatial patterning was also observed for a mixture of Cy5-22 nt RNA and ATTO488-447 nt RNA (Figure S20). In contrast, mixtures of Cy5-22 nt RNA with ATTO488-80 nt RNA (Figure S21) and of ATTO488-447 nt RNA with Alexa647-967 nt RNA (Figure S22) formed a single population of vesicles with colocalized RNA, indicating that spatial patterning is strongly dependent on the relative sizes of the RNA species.

### Effect of Amphiphile Chain Length and Headgroup Structure on Vesicle Formation

While C_12_CT supports vesicle formation, shorter (C_8_CT) or longer (C_16_CT) analogs failed to promote a productive condensate-to-vesicle transition upon cysteine treatment (Figure S23). LCMS measurements confirmed that C_8_CT failed to generate diacylated cysteine amphiphiles (Di-C_8_-Cys), instead producing predominantly monoacylated species (Mono-C_8_-Cys) (Figure S24). Reducing cysteine to 0.5 equivalents relative to C_8_CT to promote diacylation did not induce RNA-enriched vesicle formation and instead produced irregularly shaped vesicles lacking RNA (Figure S25).^44^ In contrast, C_16_CT produced diacylated cysteine (Di-C_16_-Cys) comparable to that of C_12_CT (Di-C_12_-Cys) (Figure S24). However, only aggregated RNA-lipid particles were observed, as indicated by colocalized Quantifluor and DiI signals (Figure S23). This might be due to the significantly higher gel-liquid phase transition temperature of Di-C_16_-Cys (60.4 °C)^65^ compared to Di-C_12_-Cys (30.8 °C; Figure S26), which inhibits membrane fluidity.

To test the generality of the vesicle-forming reaction, alternative reactive headgroups were tested. Cysteine was replaced with the N-terminal cysteine-containing dipeptide cysteinylglycine (Cys–Gly). Upon addition of Cys-Gly to 80 nt RNA-C_12_CT condensates, RNA-enriched vesicles formed within 1 h. These vesicles exhibited smaller diameters (0.76 ± 0.32 μm) than those generated with cysteine (Figure S27). LCMS analysis confirmed formation of the corresponding diacylated product (Di-C_12_-Cys-Gly) (Figures S27).

### RNA-Enrichment in Vesicles Enables Functional RNA Chemistry

Ribozyme catalysis is intrinsically concentration dependent as low RNA concentrations limit molecular encounter rates and weaken ribozyme substrate binding, whereas high local concentrations enhance productive interactions. Mechanisms capable of concentrating RNA within closed compartments may therefore play a critical role in enabling catalytic RNA chemistry.^66^ To test whether the increase in RNA concentration during the condensate-to-vesicle transition can activate ribozyme function, we designed a model ribozyme-substrate system and compared catalytic activity in bulk solution, RNA-C_12_CT condensates, and RNA-enriched vesicles. We employed a hairpin ribozyme (R53) derived from the tobacco ringspot virus together with its 32 nt substrate RNA (S32), which undergoes site-specific cleavage to yield 22 nt and 10 nt fragments F22 and F10 (Figure 4a).^67–69^ Initial attempts to activate ribozyme catalysis by direct addition of Mg^2+^ after vesicle formation resulted in extensive membrane rupture (Figure S28). This instability likely arises from strong interactions between Mg^2+^ and the carboxylate headgroups of the diacylated cysteine lipids. Such incompatibility between divalent cations and fatty acid-derived membranes represents a longstanding challenge in RNA protocell studies, since ribozyme catalysis typically requires millimolar Mg^2+^ concentrations.^70^ To introduce Mg^2+^ without compromising membrane integrity, we exploited the ability of condensates to recruit high local concentrations of divalent cations.^40^ Ribozyme and substrate RNA were premixed with Mg^2+^ (78 nM ribozyme, 1.3 μM substrate, and 0.1 mM Mg^2+^) and subsequently combined with C_12_CT to form RNA-C_12_CT condensates. To maintain electrostatic balance in the presence of Mg^2+^, the C_12_CT concentration was reduced to 5% of the original condition. Upon cysteine-triggered diacylation, these condensates transformed into vesicles within 20 min (Figure S29). Notably, RNA mixed with 5% C_12_CT in the absence of Mg^2+^ failed to form condensates or vesicles (Figure S8), highlighting the synergistic contribution of Mg^2+^ in maintaining charge balance. Quantifluor fluorescence intensity measurements were used to estimate a 260-fold enrichment of ribozyme and substrate RNA within vesicles, corresponding to internal concentrations of 20 μM and 340 μM, respectively (Figure S29). This enrichment exceeded that observed for 80 nt RNA-enriched vesicles and is likely facilitated by Mg^2+^-mediated RNA association. Under these conditions, ribozyme activity was strongly enhanced. A cleaved fraction of 0.4 ± 0.04 was reached after 2 h, compared with 0.03 ± 0.1 in bulk solution (Figure 4b,c, S30). To verify that ribozyme activity occurred within vesicles, we added RNase A (10^-5^ mg mL^-1^) and observed that encapsulated RNAs remained intact after 2 h of incubation, whereas RNAs in bulk solution were completely hydrolyzed (Figure S31). In parallel experiments, RNA-C_12_CT condensates containing Mg^2+^ also exhibited low ribozyme activity, reaching a cleaved fraction of 0.05 ± 0.1 after 2 h, despite the RNA enrichment (Figure 4b,c, S30). Based on our previous FRAP measurements, this low activity is likely due to the restricted molecular mobility within the highly viscous condensate phase, in contrast to the more fluid interior of vesicles (Figure S1).^40^ Since RNA enrichment during vesicle formation is length dependent, we also explored catalysis in a mixed system of functional RNAs (ribozymes and substrates) combined with short nonfunctional 10 nt RNAs. Fluorescence microscopy and urea–PAGE confirmed selective enrichment of the functional RNAs within vesicles, with exclusion of the short nonfunctional RNAs during the condensate-to-vesicle transition (Figure S32). Selective partitioning from condensates containing both functional and nonfunctional RNAs biases the composition of compartments towards longer catalytic species, further highlighting the functional advantage of selective entrapment within vesicles.

**Figure 4.**
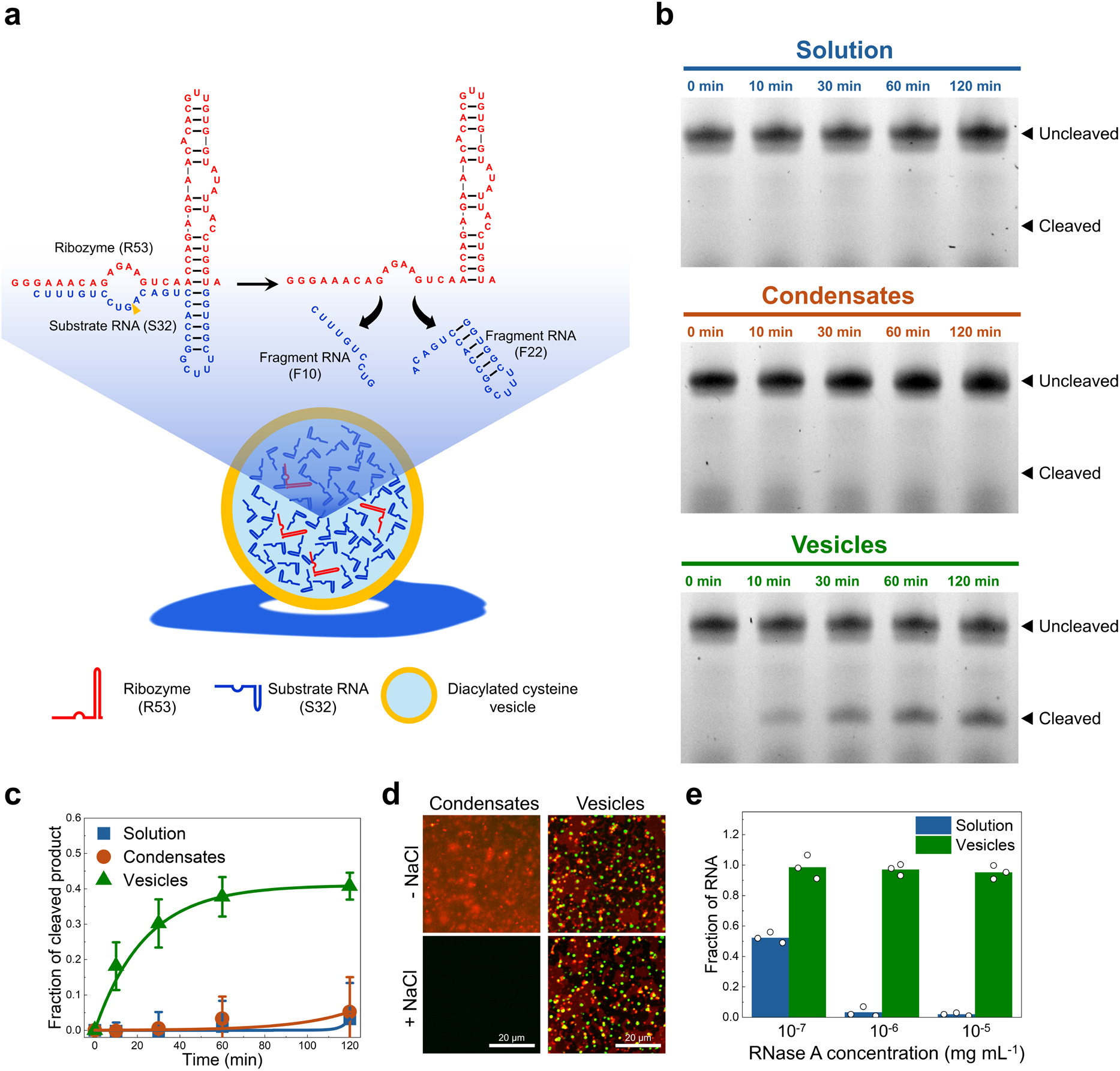
Reaction driven RNA-enrichment enables catalytic function. a) Schematic of the R53 hairpin ribozyme cleaving the 32 nt substrate (S32) to generate 22 nt (F22) and 10 nt (F10) fragments. b) Substrate cleavage by the hairpin ribozyme in bulk solution, RNA-C_12_CT condensates, and RNA-enriched vesicles. Reactions were carried out in 20 mM HEPES buffer (pH 8.2) containing 0.1 mM MgCl_2_, with final concentrations of 1.3 µM substrate and 78 nM hairpin ribozyme. Time points were taken at 0, 10, 30, 60, and 120 min. c) Fraction of cleaved product as a function of time. Data were fit to Eq. (1). Error bars indicate s.d. from three independent experiments (n = 3). Uncropped gel images are provided in Figure S30. d) Confocal fluorescence images of RNA-C_12_CT condensates and RNA-enriched vesicles before and 10 min after addition of 400 mM NaCl. Experiments used 80 nt RNA. Quantifluor (green) and DiI (red) denote RNA and membrane signals, respectively. e) Fraction of 80 nt RNA remaining after 10 min of incubation with RNase A concentrations of 10^-7^-10^-5^ mg mL^-1^ (10-fold increments) either in solution or within RNA-enriched vesicles. Points represent independent experiments (n = 3). Corresponding gel images are shown in Figure S34.

In addition to enabling catalysis, RNA-enriched vesicles exhibit enhanced durability and protect encapsulated RNA from environmental challenges. A limitation of membraneless RNA condensates is their loss of integrity in high ionic strength aqueous environments,^32,36,71^ which are commonly found on Earth and are expected in extraterrestrial oceans and brines.^72,73^ We determined the stability of RNA-C_12_CT condensates and the RNA-enriched vesicles under high-salt conditions. After 10 min of exposure to 400 mM NaCl, the RNA-C_12_CT condensates fully dissolved, whereas the RNA-enriched vesicles remained intact and retained their encapsulated RNA (Figure 4d). Osmolyte addition induced osmotic shrinkage and a reduction in vesicle size, followed by transient pearling and recovery to spherical morphologies within 30 s (Figure S33).^74,75^ Resistance to biochemical degradation was assessed by exposing free RNA and RNA-enriched vesicles to RNase A. As expected, after 10 min of treatment, free RNA in solution was 50% degraded when RNase A concentration was 10^-7^ mg mL^-1^ and nearly completely hydrolyzed when the RNase A concentration was increased to above 10^-6^ mg mL^-1^ (Figure 4e, S34). In contrast, RNA within vesicles remained largely intact (>95%) across RNase A concentrations from 10^-7^ to 10^-5^ mg mL^-1^.

## Discussion

In this work, we show that simple electrostatic interactions between RNA and cationic thioesters generate condensate intermediates that, upon chemical transformation, convert into unilamellar, RNA-enriched vesicles. This reaction-driven condensate-to-vesicle transition concentrates RNA from dilute solution, selects for specific RNA species based on length, and enables catalytic activity within membrane-bound compartments. Notably, RNA enrichment was not observed when vesicles were formed from preassembled lipid films, indicating that passive lipid self-assembly is insufficient for RNA capture. Instead, enrichment requires lipid formation during the condensate-to-vesicle transition, suggesting that in situ lipid synthesis may play an important role in coupling compartment assembly to molecular concentration in vesicular systems. Beyond its implications for the formation of protocells, the ability of this reaction-driven process to concentrate and encapsulate RNA from dilute solution may also be relevant to many other applications where efficient RNA compartmentalization is desirable.^66^

Compartment formation acts not only as a mechanism of encapsulation, but also as a form of molecular selection. Chemically gated vesicle formation intrinsically discriminates RNA encapsulation by polymer length, selectively retaining longer RNAs while shorter RNAs escape. RNA length influences both encapsulation efficiency and retention. The result is selective enrichment of longer RNA polymers while shorter oligonucleotides are excluded. It is likely that the greater multivalency of longer RNAs stabilizes condensate intermediates and slows RNA escape as thioester is consumed during membrane formation. The observed length-dependent compartmentalization provides a physical route for enriching RNAs that are more capable of folding, catalysis, and information storage, thereby linking molecular information to emergent compartment properties.^39,65^ RNA length appears to act as a variable influencing vesicle size, stability, and function, thus establishing a primitive genotype–phenotype coupling in which informational content biases the physical characteristics of the resulting membrane-bound compartments.

A striking result is our observation that continuous gradients of the cysteine chemical trigger result in the generation of well-defined populations of vesicles with distinct RNA compositions. Rather than exhibiting a gradual change in RNA composition, distinct vesicle populations were separated by sharp boundaries, suggesting threshold-dependent behavior during the condensate-to-vesicle transition. One rationale is that modest changes in the kinetics of membrane lipid synthesis may produce disproportionately large changes in the retention of shorter RNAs, resulting in a nonlinear response to the cysteine gradient. The multivalent interactions that influence condensate stability, membrane permeability during reaction, or phase transitions within the assembling membrane may contribute to the observed phenomenon. The conversion of a continuous chemical gradient into discrete compartment populations is reminiscent of the threshold patterning observed in morphogen-driven developmental systems. However, here the effect arises entirely from physicochemical interactions rather than genetic regulation.

The emergence of distinct compartment populations from a continuous chemical gradient is significant because it demonstrates a physicochemical mechanism by which an initially homogeneous molecular mixture can spontaneously generate spatially differentiated states. Living systems are characterized not only by compartmentalization, but also by the coexistence of chemically distinct populations of cells that have different functional roles. Cell populations differ not merely in their location but also in the informational polymers that they contain. Because RNA length strongly influences folding, molecular recognition, catalytic activity, and information storage, gradient-directed compartment formation has the potential to generate compartments with distinct functional potential from a common starting mixture. In the absence of pre-existing biological machinery, environmental chemical gradients provide a route for generating spatially organized molecular diversity, offering a possible mechanism by which chemical environments can drive differentiation and functional heterogeneity during the transition from non-living matter to living systems.

The activation of ribozyme catalysis following the condensate-to-vesicle transition highlights a functional consequence of reaction-driven compartment formation. Many biological processes require local concentrations exceeding a critical threshold required for catalysis or molecular recognition. By concentrating both ribozyme and substrate within membrane-bound compartments, reaction drives the formation of catalytically active microenvironments from otherwise inactive dilute solutions.

The condensate-to-vesicle pathway also benefits from the presence of Mg^2+^, a cofactor that is essential for many RNA-catalyzed processes and central to contemporary biochemistry. In contrast to fatty-acid-based protocell systems, where Mg^2+^ frequently destabilizes membranes or promotes aggregation, here Mg^2+^ facilitates compartment formation by stabilizing condensate formation and allowing vesicle formation using substantially lower concentrations of the cationic thioester. The condensate intermediates may also be concentrating Mg^2+^ alongside RNA, creating local environments that are favorable for ribozyme activity.

The condensate-to-vesicle mechanism described here couples chemical reactions to the selective concentration of RNA, converting initially dilute molecular mixtures into functional compartments. These findings illustrate how reaction-driven compartmentalization can simultaneously generate molecular selection, spatial organization, and catalytic activity, linking simple chemical processes to key features of living systems.

