## Supplementary Information for "A Reaction-Driven Condensate-to-Vesicle Transition Selects, Activates, and Spatially Organizes RNA"

**Materials and general methods**

All reagents and solvents employed were commercially available and used as supplied without further purification. Quantifluor RNA dye was purchased from Promega, and DiI (1,1'-dioctadecyl-3,3,3',3'-tetramethylindocarbocyanine perchlorate) was purchased from Thermo Fisher Scientific. Choline thioester derivatives, including $C_8$CT, $C_{12}$CT, and $C_{16}$CT were synthesized as previously described.[1,2] tRNA from baker's yeast was purchased from Merck. tRNA used in this study was assumed to have a length of 80 nt, according to the supplier's specification (Merck, product R5636). Because the exact sequence composition was not provided, molecular weight calculations were performed assuming an average nucleotide mass of 330 Da. Based on this approximation, the molecular weight of the 80 nt tRNA was estimated to be ~26.4 kDa. The 10, 22, 447, 967 and 976 nt RNAs, together with the hairpin ribozyme and substrate RNA, were either prepared using a T7 transcription kit or purchased from IDT or Cisterna, as summarized in Table 1. All confocal microscopy was conducted using 8-well glass bottom chambers (Ibidi) on a W1-SoRA spinning disk confocal microscope system (Nikon) equipped with 60 x oil-immersion objective lens using a combination of 488 nm laser and FITC filter set (for Quantifluor and ATTO488), 561 nm laser and TRITC filter set (for DiI), and 640 nm laser and Cy5 filter set (for Cy5 and Alexa647). Super-resolution imaging was performed using a SORA (spinning-disk confocal super-resolution by optical pixel reassignment) unit mounted on the confocal microscopy system. All images were digitally developed using image J software.

**Preparation of Fluorescently labeled RNAs.** Cy5-labeled 10 nt and 22 nt RNAs were obtained by custom synthesis from IDT. ATTO488-labeled 80 nt and 447 nt RNAs were prepared from unlabeled 80 nt RNA (purchased from Merck) and 447 nt RNA (custom-ordered from Cisterna Biologics) using the 3' End Labeling of RNA with T4 RNA Ligase kit (New England Biolabs) and pCp-ATTO488 (Gena Bioscience), according to the manufacturer's instructions. Labeled RNAs were subsequently purified from the reaction mixture using a Monarch Spin RNA Cleanup Kit (New England Biolabs).

Alexa647-labeled 967 nt RNA (mCherry mRNA) was prepared as follows. Azide-modified mRNA was transcribed using the TranscriptAid T7 High Yield Transcription Kit (Thermo Fisher Scientific) according to the manufacturer's instructions. Briefly, a plasmid encoding mCherry under a T7 promoter was linearized by XbaI digestion. The transcription reaction contained 1 µg linearized template, 10 mM ATP, 10 mM CTP, 10 mM GTP, 7.5 mM UTP and 2.5 mM 5-Azido-C3-UTP, and was incubated at 37 °C for 2 h. After DNase treatment, the mRNA was recovered by ethanol precipitation. The RNA pellet was washed, dried and resuspended in TE buffer. Purified azide-modified mRNA (500 ng $\mu L^{-1}$) was then reacted with AF647-DBCO at a final concentration of 1 mM and incubated at 37 °C for 2 h. Excess AF647-DBCO was subsequently removed using a Monarch Spin RNA Cleanup Kit (New England Biolabs) according to the manufacturer's instructions.

**Confocal microscopy of RNA-$C_{12}$CT condensates and in-situ vesicle formation.** RNA solution (100 µL, 0.1 mg $mL^{-1}$ in 60 mM sodium phosphate buffer, pH 8.2) was placed inside a single well of an 8-well glass bottom chamber slide. Unless otherwise specified, all RNAs used in this manuscript were prepared at the same volume and concentration. For experiments involving mixtures of two RNA species, including 10 nt RNA with 80 nt RNA, Cy5-22 nt RNA with 976 nt RNA, Cy5-22 nt RNA with ATTO488-80 nt RNA, Cy5-22 nt RNA with ATTO488-447 nt RNA, and ATTO488-447 nt RNA with Alexa647-967 nt RNA,

1:1 (w/w) RNA mixtures were prepared to give a final RNA solution of 100 μL, 0.1 mg $mL^{-1}$ in 60 mM sodium phosphate buffer (pH 8.2). In the general procedure, Quantifluor (0.2 μL of the original stock solution) and DiI (0.2 μL, 1 mM in MeOH) solutions were added into the RNA solution. For experiments using ATTO488-labeled RNAs, Quantifluor was omitted. $C_{12}CT$ solution (100 μL, 12 mM in DIW) was added to the RNA solution to form RNA-$C_{12}CT$ condensates. For the experiment using CT derivatives, the same concentration of $C_8CT$ and $C_{16}CT$ was used instead of $C_{12}CT$. After incubation at 37 °C for 5 min, cysteine solution (100 μL, 30 mM in DIW) was added to the RNA-$C_{12}CT$ condensates and homogenized by pipette mixing, followed by further incubation at 37 °C. For spatial patterning experiments, the same cysteine solution was instead introduced as a single, gentle addition at the far-left edge of the well, without any subsequent mixing or agitation. The RNA-$C_{12}CT$ condensates at the bottom of the chamber and their changes upon the cysteine treatment were visualized by confocal fluorescence microscopy (Nikon).

**Composition index (CI) analysis.** To quantify the relative composition of short and long RNAs within individual vesicles, we defined a composition index (CI) based on normalized fluorescence intensities. For each RNA pair, the fluorescence signal corresponding to the short RNA was normalized by setting the mean fluorescence intensity of vesicles located within 50 μm of the left edge of the image to 1 and the background fluorescence to 0. Likewise, the fluorescence signal corresponding to the long RNA was normalized by setting the mean fluorescence intensity of vesicles located within 50 μm of the right edge of the image to 1 and the background fluorescence to 0. The CI for each vesicle was then calculated as

$$\mathrm{CI} = \frac{I_{\mathrm{short}} - I_{\mathrm{long}}}{I_{\mathrm{short}} + I_{\mathrm{long}}}$$

where $I_{\mathrm{short}}$ and $I_{\mathrm{long}}$ are the normalized fluorescence intensities of the short and long RNAs, respectively, measured for each individual vesicle.

**Confocal microscopy of RNA-thioester condensate intermediates.** 80 nt RNA solution (100 µL, 0.1 mg mL$^{-1}$ in 60 mM sodium phosphate buffer, pH 8.2) was placed inside a single well of an 8-well glass bottom chamber slide. Quantifluor (0.2 µL of the original stock solution) and DiI (0.2 µL, 1 mM in MeOH) solutions were added into the RNA solution. $C_{12}$CT solution (100 µL, 12 mM in DIW) was added to the RNA solution to form RNA-$C_{12}$CT condensates. After incubation at 37 °C for 5 min, cysteine solution (100 µL, 1.2, 2.4, 3.6 or 4.8 mM in DIW, corresponding to 0.1, 0.2, 0.3 or 0.4 eq. relative to $C_{12}$CT) was added to the RNA-$C_{12}$CT condensates and homogenized by pipette mixing, followed by further incubation at 37 °C. After overnight incubation, samples were imaged by confocal fluorescence microscopy (Nikon).

**Harpin ribozyme assay.** Ribozyme R53 (0.4 µL, 1 µg µL$^{-1}$ in DIW) and substrate S32 (4 µL, 1 µg µL$^{-1}$ in DIW) were combined and denatured at 96 °C for 5 min, and then cooled to 20 °C. The RNA mixture was then diluted into buffer (100 µL, 60 mM HEPES, 0.3 mM $MgCl_2$, pH 8.2) and transferred to an 8-well glass-bottom chamber. To generate RNA-$C_{12}$CT condensates, $C_{12}$CT (100 µL, 0.6 mM in DIW) was added and the mixture was incubated at 37 °C for 10 min. For the solution control, an equal volume of DIW was added in place of $C_{12}$CT. Vesicle formation was induced by addition of cysteine (100 µL, 30 mM in DIW). For the solution and condensate controls, DIW was added instead of cysteine.

Reactions were incubated at 37 °C, and aliquots were collected at 0, 10, 30, 60, and 120 min. The resulting mixtures were harvested after scratching the bottom of the chamber with a disposable spatula and subjected to 15% Urea-PAGE. The electrophoresis was conducted at 200 V for 50 min followed by 5 min of Gel red staining. The resulting gels were visualized by Gel imaging (Bio-Rad). The relative amount of substrates and cleaved products on the gel was estimated by using Image Lab software (Bio-Rad). The fraction of the cleaved product was calculated by using the equation A/(A+B), where A is the sum of intensities for cleaved bands, and B is the intensity for unreacted substrates. For kinetic experiments, data were fit to a first order exponential

$$f(t) = f_{max}(1 - e^{(-k_{obs} \cdot t)}) \quad (1)$$

Where f(t) is the fraction product formed at indicated time t, $f_{max}$ is the plateau, and $k_{obs}$ is the observed rate constant. All fits were performed in Origin 2025.

**RNA-$C_{12}$CT condensates and RNA enriched vesicles in high salt conditions.** 80 nt RNA solution (100 μL, 0.1 mg $mL^{-1}$ in 60 mM sodium phosphate buffer, pH 8.2) was placed in a well of an 8-well glass-bottom chamber slide. Quantifluor (0.2 μL of the stock solution) and DiI (0.2 μL, 1 mM in MeOH) were added to the RNA solution. $C_{12}$CT solution (100 μL, 12 mM in DIW) was added to the RNA solution to form RNA-$C_{12}$CT condensates. After 5 min of incubation at 37 °C, a cysteine solution (100 μL, 30 mM in DIW) was added to the RNA-$C_{12}$CT condensates, which were then further incubated at 37 °C. For the RNA-$C_{12}$CT condensate control, the same volume of DIW was added instead of cysteine solution. After 10 min of incubation at 37 °C, NaCl solution (3 μL, 4 M in DIW) was added to both RNA-$C_{12}$CT condensates and RNA-enriched vesicles, resulting in a final NaCl concentration of 400 mM. The resulting RNA-$C_{12}CT$ condensates and RNA-enriched vesicles were imaged using confocal fluorescence microscopy (Nikon).

**RNA hydrolysis assay.** The 80 nt RNA solution (100 μL, 0.1 mg $mL^{-1}$ in 60 mM sodium phosphate buffer, pH 8.2) was placed inside a well of an 8-well glass bottom chamber. $C_{12}CT$ solution (100 μL, 12 mM in DIW) was added to the RNA solution to form RNA-$C_{12}CT$ condensates. After 5 min of incubation at 37 °C, cysteine solution (100 μL, 30 mM in DIW) was added to the RNA-$C_{12}CT$ condensates and the resulting mixture was incubated for 10 min at 37 °C. For the experiment varying with RNase A concentration, various RNase A solutions (0.3 μL, $10^{-2}$, $10^{-3}$, and $10^{-4}$ mg $mL^{-1}$) were added to the RNA, $C_{12}CT$ and cysteine mixtures to give final RNase A concentration of $10^{-5}$, $10^{-6}$, and $10^{-7}$ mg $mL^{-1}$, respectively. After incubation at 37 °C for 10 min, RNase inhibitor (10 μL, 20 U $\mu L^{-1}$) was added to each mixture. For the solution control, all conditions were kept the same, but an equal volume of DIW was added instead of the $C_{12}CT$ solution. The resulted mixtures were harvested after scratching the bottom of the chamber with a disposable spatula and were subjected to the 5% Urea-PAGE. The electrophoresis was conducted at 200 V for 20 min followed by 5 min of Gel red staining. The resulting gels were visualized by Gel imaging (Bio-Rad). The relative amount of RNA on the gel was estimated by using Image Lab software (Bio-Rad).

**Table 1.** Information on RNAs

| Name | Sequence | Source | Notes |
|---|---|---|---|
| 10 nt RNA | 5'-CAGUACAAUC-3' | IDT | Fragment of T7p14 vector, Cy5 labeled on 5' |
| 15 nt RNA | 5'- AGAACUGCUAACUCA-3' | IDT | tRF-2 |
| 22 nt RNA | 5'-UAGCUUAUCAGACUGAUGUUGA-3' | IDT | miR-21, Cy5 labeled on 5' |
| Ribozyme (53 nt) | 5'-<br>GGGAAACAGAGAAGUCAACCAGAGAAACACACGUUGU<br>GGUAUAUUACCUGGUA-3' | Cisterna | Fragment of R3C ligase |
| Substrate RNA (32 nt) | 5'-GGUGGCUUUCGGCCACCUGACAGUCCUGUUUC-3' | Cisterna | Fragment of R3C ligase |
| 447 nt RNA | 5'-<br>GUUUAACUUUAAGAAGGAGAUAUACCGGGAACCCCC<br>CUUCGGGGGGUCACCUCGCGCAGCGGGCUGCGCGA<br>AGGAGCCACGCUGCGAAGCAGCGUGGCGGUUCUCG<br>UGCGUCACCGAAACGCACGAAGGUCGCGCCUCUUCA<br>CGAGGCGUCACCUGGGAGAGCGCGAAAGCGCUAGCC<br>CGUGCUCUAGCAGGCCUCGAGAUCUCCUCUAGAAAU<br>AAUUUUGUUUAACUUUAAGAAGGAGAUAUACAUAACU<br>UGGUCCGAAUACGGGUCCUAGCAACGUUCGGGCACG<br>UACGAAGGAAGGUUUGGUAUGUGGUAUAUUCGUACG<br>UGCCGGUACCUCUAGAGUCGACCCGGGCGGCCGCU<br>UGCGGCCGCACUCGAGAGAUCUAGAGCGUCACGGUC | Cisterna | UMDV-MangoIII-DMDV, ATTO488 labeled on 3' |

| | | | |
|---|---|---|---|
| | GAACUCCCGUACGAGGUGCCCGCACCUCGUCCCCCC CUUCCGGGGGGGUCCCC-3' | | |
| 967 nt RNA | 5'- GGACUCUUCUGGUCCCCACAGACUCAGAGAGAACCC ACCAUGGUGAGCAAGGGCGAGGAGGAUAACAUGGCC AUCAUCAAGGAGUUCAUGCGCUUCAAGGUGCACAUG GAGGGCUCCGUGAACGGCCACGAGUUCGAGAUCGAG GGCGAGGGCGAGGGCCGCCCCUACGAGGGCACCCA GACCGCCAAGCUGAAGGUGACCAAGGGUGGCCCCCU GCCCUUCGCCUGGGACAUCCUGUCCCCUCAGUUCAU GUACGGCUCCAAGGCCUACGUGAAGCACCCCGCCGA CAUCCCCGACUACUUGAAGCUGUCCUUCCCCGAGGG CUUCAAGUGGGAGCGCGUGAUGAACUUCGAGGACGG CGGCGUGGUGACCGUGACCCAGGACUCCUCCCUGCA GGACGGCGAGUUCAUCUACAAGGUGAAGCUGCGCGG CACCAACUUCCCCUCCGACGGCCCCGUAAUGCAGAA GAAGACCAUGGGCUGGGAGGCCUCCUCCGAGCGGAU GUACCCCGAGGACGGCGCCCUGAAGGGCGAGAUCAA GCAGAGGCUGAAGCUGAAGGACGGCGGCCACUACGA CGCUGAGGUCAAGACCACCUACAAGGCCAAGAAGCC CGUGCAGCUGCCCGGCGCCUACAACGUCAACAUCAA GUUGGACAUCACCUCCCACAACGAGGACUACACCAU CGUGGAACAGUACGAACGCGCCGAGGGCCGCCACUC CACCGGCGGCAUGGACGAGCUGUACAAGUAGCUCGA GGCUGGAGCCUCGGUGGCCAUGCUUCUUGCCCCUU GGGCCUCCCCCCAGCCCCUCCUCCCCUUCCUGCACC | Using T7 transcription kit | Alexa647-labeled 967 nt RNA was prepared by coupling Alexa647-DBCO to azide-modified mCherry RNA. |

| | | | |
|---|---|---|---|
| | CGUACCCCCGUGGUCUUUGAAUAAAGUCUGAGUGGG<br>CGGCAAAAAAAAAAAAAAAAAAAAAAAAAAAAAAAAAAA<br>AAAAAAAAAAAAAAAAAAAAAAAAAAAAAAAAAAAAAAA<br>AAAAAAAAAAAAAAAAAAAAAAAAAA-3’ | | |
| 976 nt RNA | 5'-<br>GGACUCUUCUGGUCCCCACAGACUCAGAGAGAACCC<br>ACCAUGGUGAGCAAGGGCGAGGAGCUGUUCACCGGG<br>GUGGUGCCCAUCCUGGUCGAGCUGGACGGCGACGU<br>AAACGGCCACAAGUUCAGCGUGUCCGGCGAGGGCGA<br>GGGCGAUGCCACCUACGGCAAGCUGACCCUGAAGUU<br>CAUCUGCACCACCGGCAAGCUGCCCGUGCCCUGGCC<br>CACCCUCGUGACCACCCUGACCUACGGCGUGCAGUG<br>CUUCAGCCGCUACCCCGACCACAUGAAGCAGCACGA<br>CUUCUUCAAGUCCGCCAUGCCCGAAGGCUACGUCCA<br>GGAGCGCACCAUCUUCUUCAAGGACGACGGCAACUA<br>CAAGACCCGCGCCGAGGUGAAGUUCGAGGGCGACAC<br>CCUGGUGAACCGCAUCGAGCUGAAGGGCAUCGACUU<br>CAAGGAGGACGGCAACAUCCUGGGGCACAAGCUGGA<br>GUACAACUACAACAGCCACAACGUCUAUAUCAUGGCC<br>GACAAGCAGAAGAACGGCAUCAAGGUGAACUUCAAGA<br>UCCGCCACAACAUCGAGGACGGCAGCGUGCAGCUCG<br>CCGACCACUACCAGCAGAACACCCCCAUCGGCGACG<br>GCCCCGUGCUGCUGCCCGACAACCACUACCUGAGCA<br>CCCAGUCCGCCCUGAGCAAAGACCCCAACGAGAAGC<br>GCGAUCACAUGGUCCUGCUGGAGUUCGUGACCGCCG<br>CCGGGAUCACUCUCGGCAUGGACGAGCUGUACAAGU<br>AACUCGAGGCUGGAGCCUCGGUGGCCAUGCUUCUUG | Cisterna | eGFP |

| | |
|---|---|
| | CCCCUUGGGCCUCCCCCCAGCCCCUCCUCCCCUUCC UGCACCCGUACCCCCGUGGUCUUUGAAUAAAGUCUG AGUGGGCGGCAAAAAAAAAAAAAAAAAAAAAAAAAAAA AAAAAAAAAAAAAAAAAAAAAAAAAAAAAAAAAAAAAAA AAAAAAAAAAAAAAAAAAAAAAA-3' |

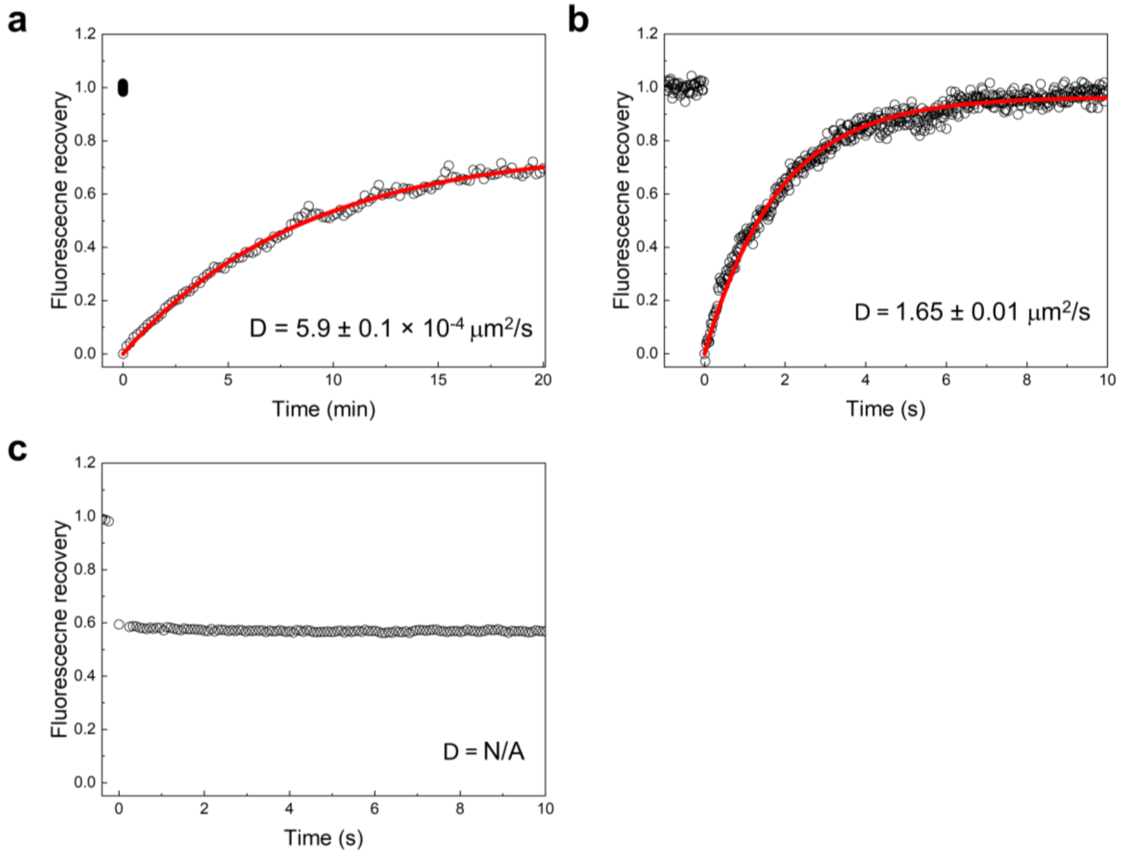

**Figure S1.** Fluorescence recovery after photobleaching (FRAP) analysis of Quantifluor signals from a) RNA-$C_{12}$CT condensates, b) RNA-thioester Condensate intermediates produced by mixing RNA-$C_{12}$CT condensates with 0.1 equivalent of cysteine relative to $C_{12}$CT, and c) the interior of vesicles. The diffusion constant (D) increased by three orders of magnitude when the RNA-$C_{12}$CT condensates transitioned into coacervate intermediates, indicating that the RNA-$C_{12}$CT condensates became more fluid. The D value inside the vesicles could not be measured, as the diffusion rate was faster than the acquisition speed (10 ms). D values were calculated as previously described.[3] All experiments were performed using 80 nt RNA.

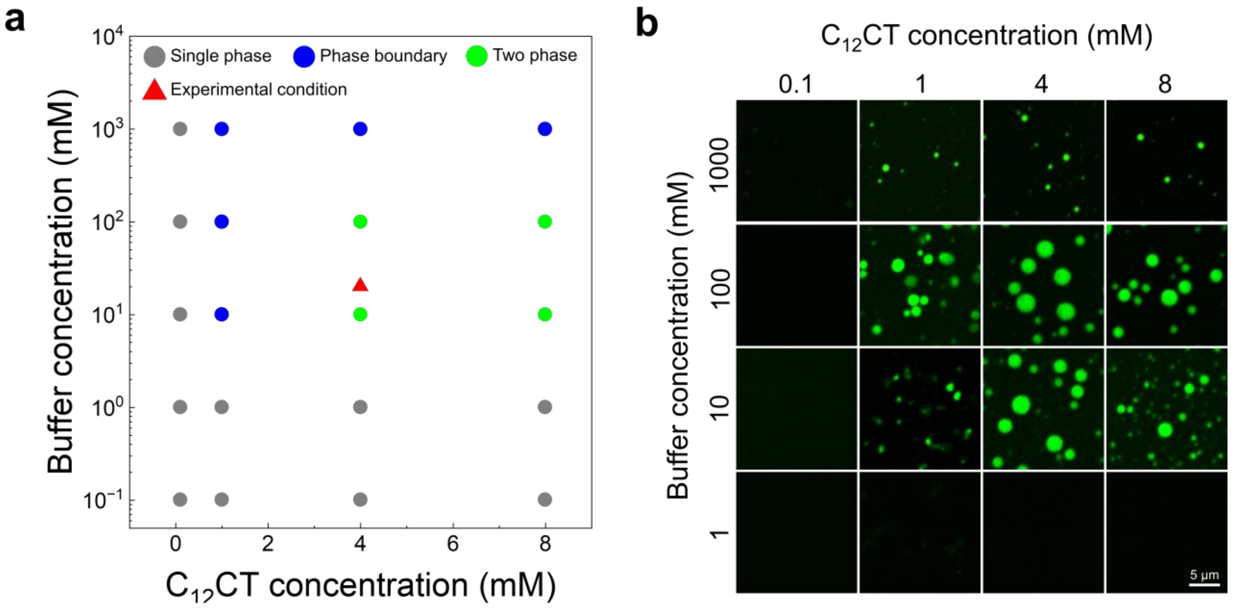

**Figure S2.** a) Phase diagram of the mixture of 80 nt RNA and $C_{12}CT$ as a function of $C_{12}CT$ concentration and sodium phosphate buffer concentration. The RNA concentration was maintained at 1.3 µM. Gray circles indicate the single-phase region, blue circles denote the phase boundary corresponding to the onset of condensate formation, and green circles represent the two-phase region containing abundant RNA-$C_{12}CT$ condensates. Red triangles mark the experimental conditions used to prepare RNA-enriched vesicles. b) Representative confocal fluorescence images of RNA-$C_{12}CT$ condensates across $C_{12}CT$ and buffer concentrations. RNA was visualized using Quantifluor staining.

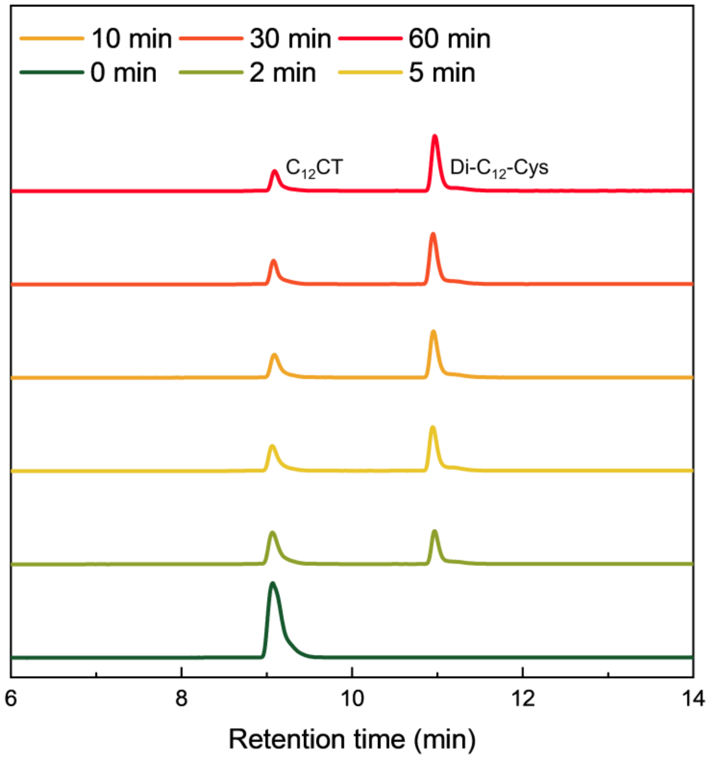

**Figure S3.** LCMS measurement of $C_{12}CT$ and diacylated cysteine (Di-$C_{12}$-Cys) after 80 nt RNA-$C_{12}CT$ condensates are treated with cysteine.

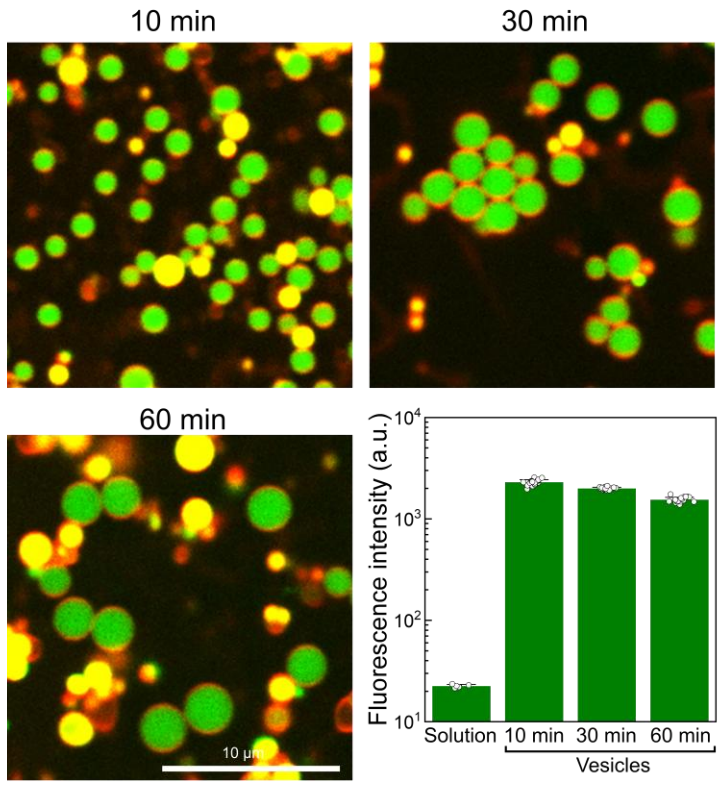

**Figure S4.** Confocal images of 80 nt RNA-enriched vesicles after 10, 30, and 60 min. Quantifluor (green) and DiI (red) denote RNA and membrane signals, respectively. The graph shows average Quantifluor intensity in bulk solution and within vesicles (n = 100). RNA fluorescence inside vesicles is enriched 102 ± 6.2 fold relative to solution at 10 min and decreases gradually to 88 ± 2.4 fold after 30 min and 68 ± 3.9 fold after 60 min (67% of the 10 min value).

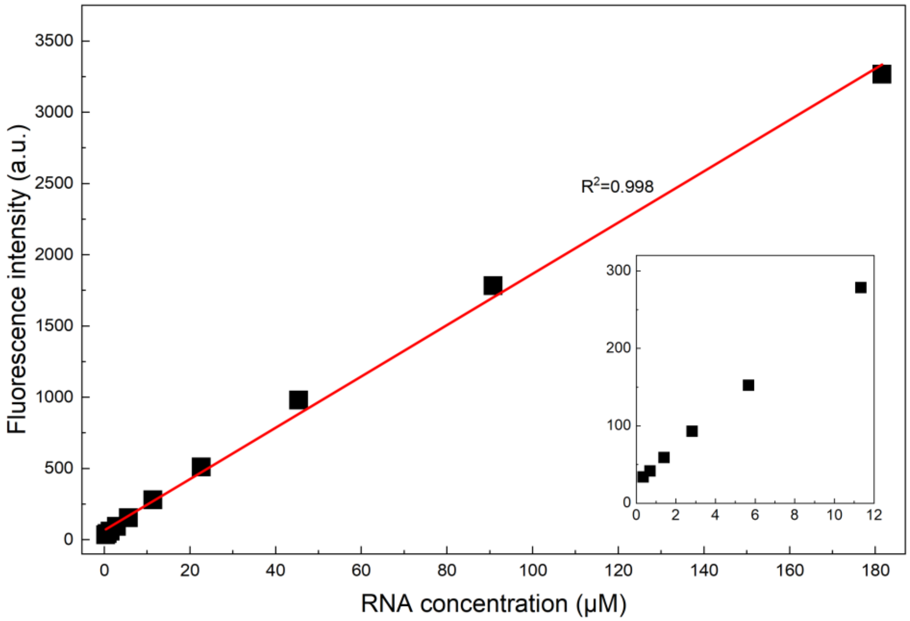

**Figure S5.** Standard curve correlating Quantifluor fluorescence intensity with RNA concentration. An 80 nt RNA stock solution (100 μL, 362 μM in 10 mM Tris buffer, pH 7.4, containing 1 mM EDTA) was mixed with Quantifluor dye (0.2 μL) in sodium phosphate buffer (100 μL, 20 mM, pH 8.2). The resulting solution was serially diluted by 50% at each step using the same sodium phosphate buffer as diluent, yielding RNA concentrations down to 0.35 μM. In the RNA-enriched vesicle system, the molar ratio of RNA to Quantifluor remains constant over time, however, Quantifluor-bound RNA becomes locally concentrated within vesicles, resulting in increased local fluorescence intensity. To mimic this condition and construct an appropriate calibration curve, Quantifluor-bound RNA was diluted into buffer lacking additional Quantifluor dye. This approach assumes that Quantifluor binding to RNA is sufficiently strong to prevent significant dissociation during serial dilution. The observed linear correlation between RNA concentration and fluorescence intensity supports the validity of this assumption. Fluorescence intensities for the standard curve samples were measured using the same confocal microscopy settings employed for imaging RNA-enriched vesicles.

**a**

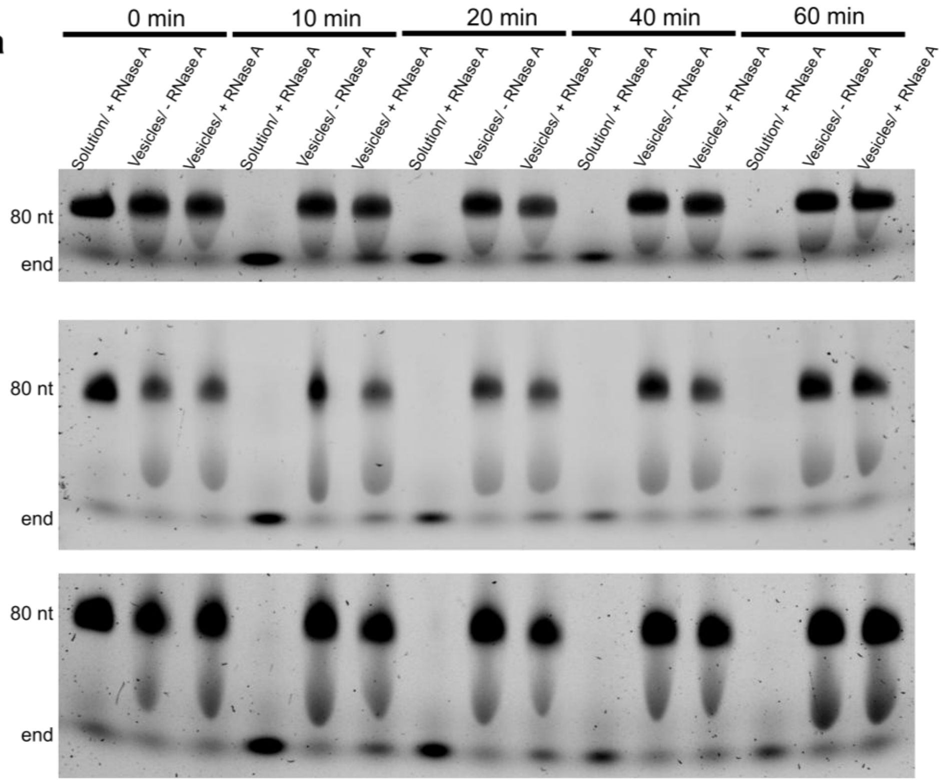
0 min
10 min
20 min
40 min
60 min
Solution/ + RNase A
Vesicles/ - RNase A
Vesicles/ + RNase A
80 nt
end
80 nt
end
80 nt
end

**b**

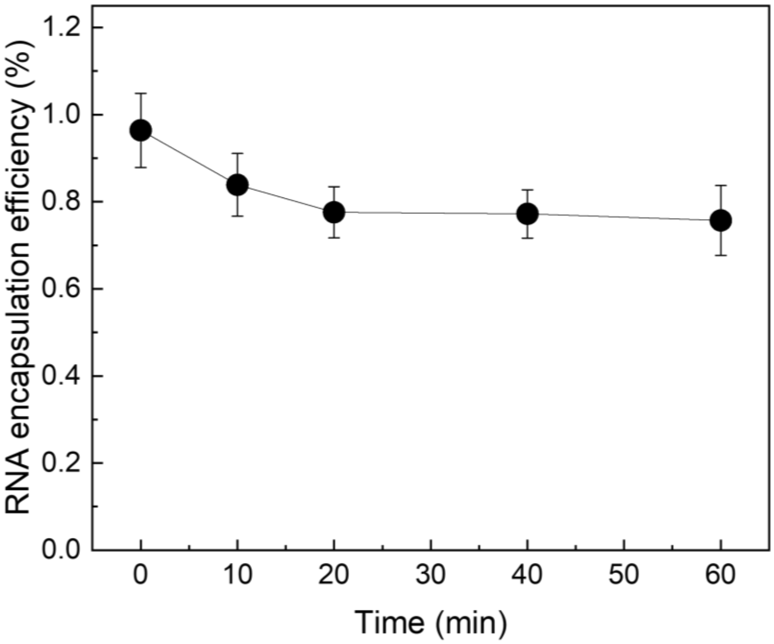
RNA encapsulation efficiency (%)
1.2
1.0
0.8
0.6
0.4
0.2
0.0
0
10
20
30
40
50
60
Time (min)

**Figure S6.** Estimation of RNA encapsulation efficiency inside the vesicles. a) Denaturing Urea-PAGE analysis of RNA solution with RNase A ($10^{-5}$ mg $mL^{-1}$), RNA-enriched vesicles without RNase A, and RNA-enriched vesicles with RNase A over 1 h of incubation (n = 3). 80 nt RNA was used in the experiment. The incubation time was measured starting 10 min after treatment of the RNA-$C_{12}$CT condensates with cysteine. Simultaneously, RNase A was added to the vesicles 10 min after the cysteine treatment. The disappearance of RNA bands in the RNase A-treated solution samples indicates that all RNAs outside the vesicles were hydrolyzed. Therefore, the remaining RNA signals in the RNase A-treated vesicle samples represent RNAs encapsulated inside the vesicles. By comparing the total RNA inside and outside of the vesicles (vesicles without RNase A) and the encapsulated RNA signals (vesicles with RNase A), the RNA encapsulation efficiency was estimated. b) Statistical evaluation of RNA encapsulation efficiency over 1 h of incubation (n = 3). The experiments were conducted at 37 ˚C.

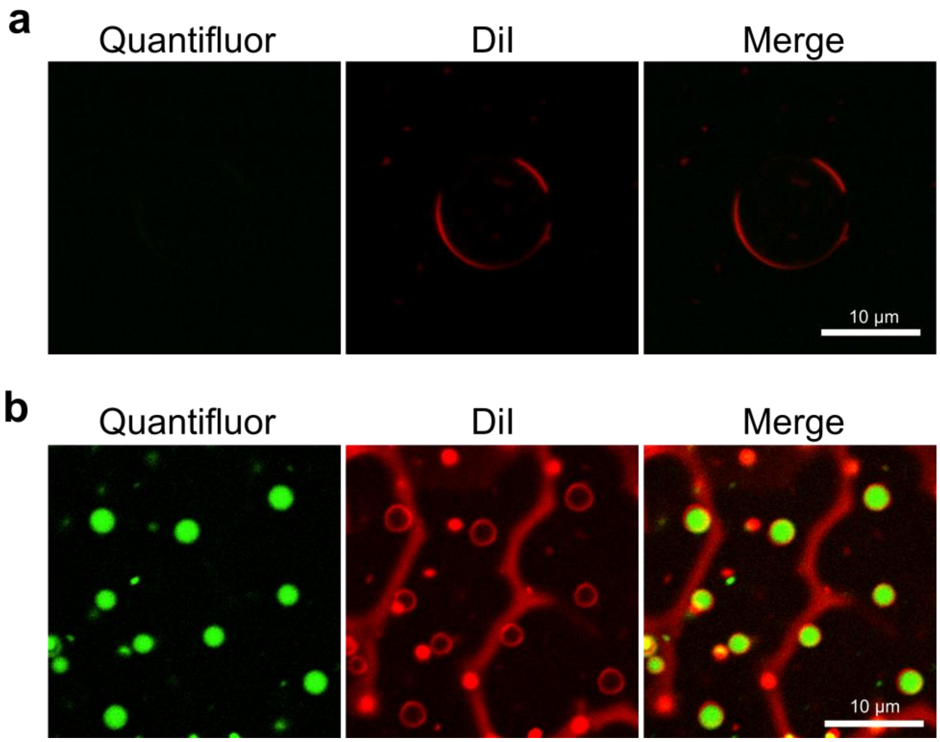

**Figure S7.** Diacylated cysteine lipid vesicles generated by a) simple hydration and b) cysteine-triggered transformation of RNA-$C_{12}$CT condensates into vesicles. Images were acquired using identical brightness and contrast settings. All experiments were performed using the same RNA solution (80 nt RNA, 0.1 mg $mL^{-1}$ in 60 mM sodium phosphate buffer, pH 8.2).

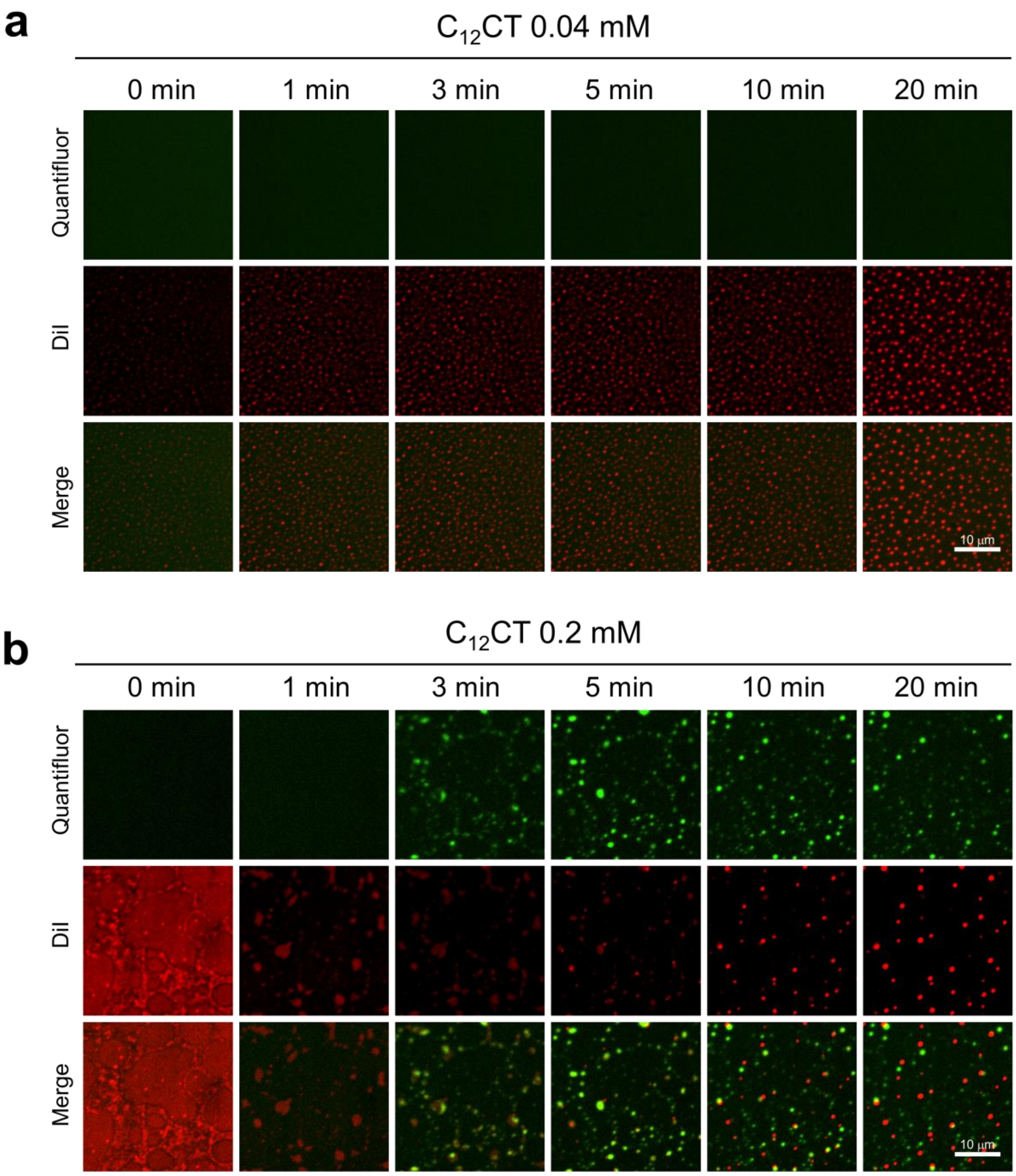
a
$C_{12}CT$ 0.04 mM
0 min
1 min
3 min
5 min
10 min
20 min
Quantifluor
DiI
Merge
10 μm
b
$C_{12}CT$ 0.2 mM
0 min
1 min
3 min
5 min
10 min
20 min
Quantifluor
DiI
Merge
10 μm

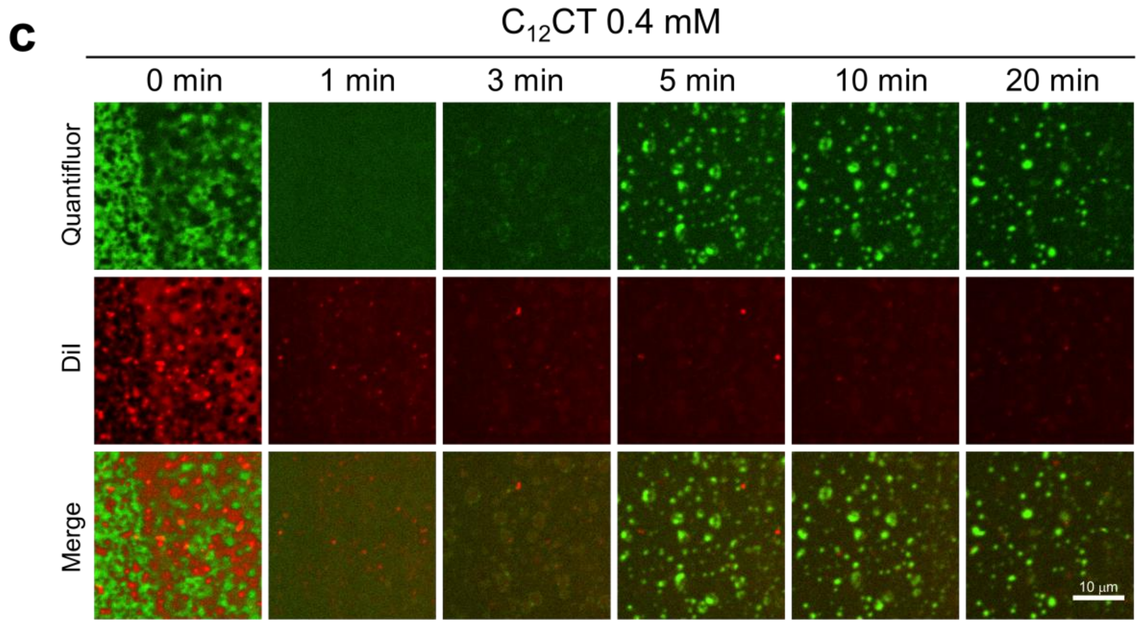

c
$C_{12}$CT 0.4 mM
0 min
1 min
3 min
5 min
10 min
20 min
Quantifluor
DiI
Merge
10 μm

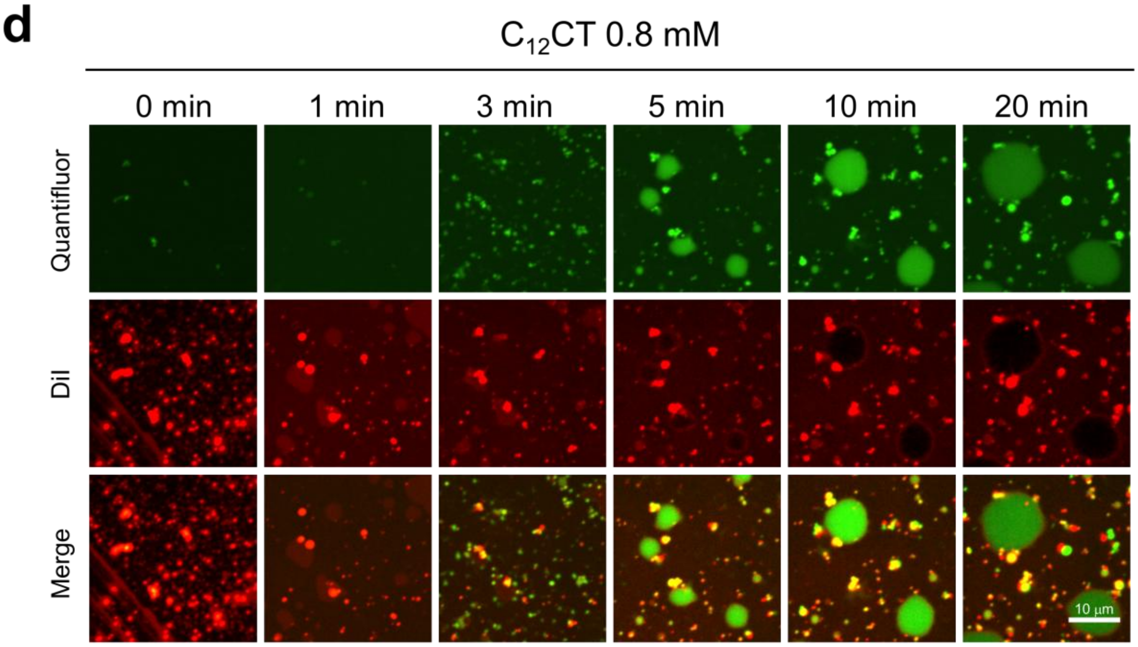

d
$C_{12}$CT 0.8 mM
0 min
1 min
3 min
5 min
10 min
20 min
Quantifluor
DiI
Merge
10 μm

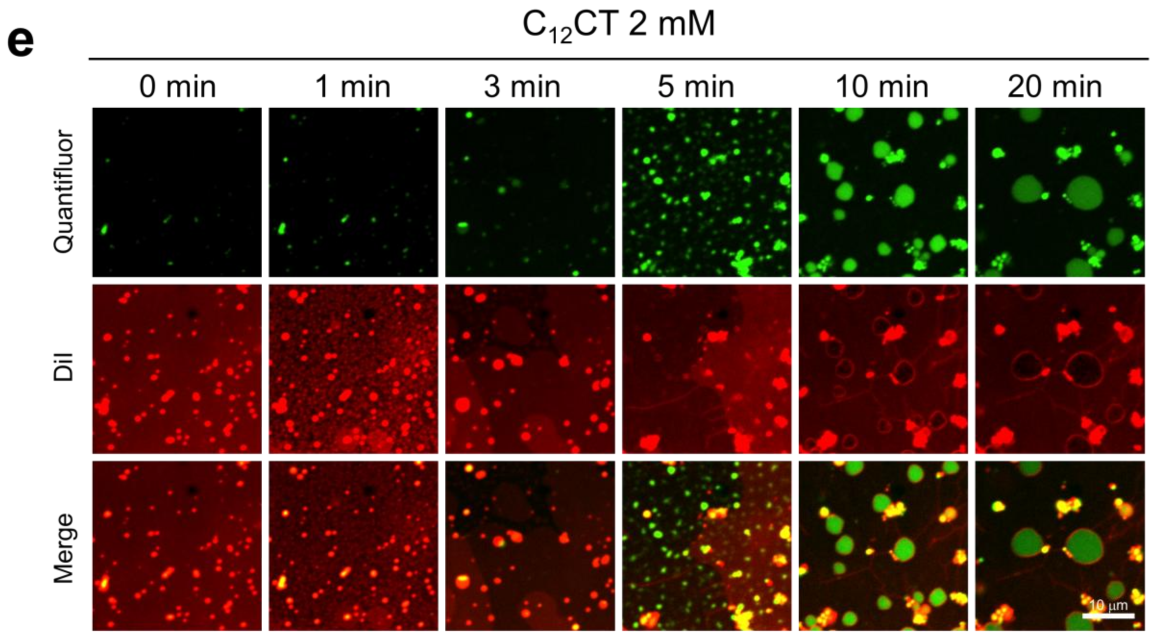

**Figure S8.** Time-lapse confocal fluorescence microscopy images of 80 nt RNA-$C_{12}$CT condensates at varying $C_{12}$CT concentrations upon cysteine treatment. $C_{12}$CT concentrations: a) 0.04 mM, b) 0.2 mM, c) 0.4 mM, d) 0.8 mM, and e) 2 mM. Relative to the standard 4 mM condition, 2 mM and 0.8 mM C12CT produced enlarged, unstable vesicles, whereas no vesicles formed at ≤0.4 mM. All experiments were performed at 37 °C.

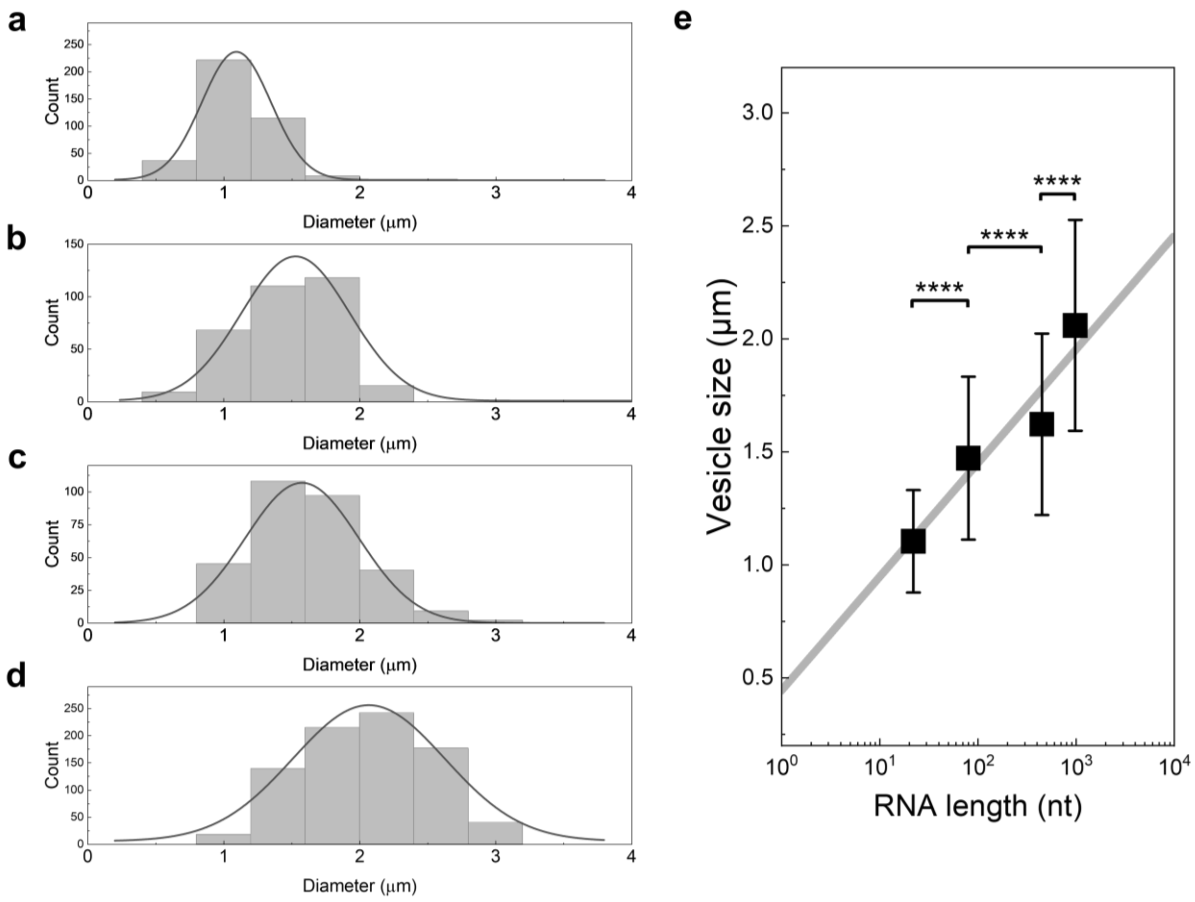

**Figure S9.** Size distributions of RNA-enriched vesicles measured 10 min after cysteine addition to the corresponding RNA-$C_{12}$CT condensates. Vesicles encapsulating (a) 22 nt, (b) 80 nt, (c) 447 nt, and (d) 976 nt RNAs were analyzed (n = 379 for 22 nt RNA, n = 320 for 80 nt RNA, n = 301 for 447 nt RNA, and n = 831 for 976 nt RNA). The mean vesicle diameters were 1.1 ± 0.25 μm for 22 nt RNA, 1.5 ± 0.36 μm for 80 nt RNA, 1.6 ± 0.43 μm for 447 nt RNA, and 2.0 ± 0.46 μm for 976 nt RNA. e) Relationship between RNA length and the diameter of RNA-enriched vesicles measured 10 min after cysteine treatment of RNA-$C_{12}$CT condensates. Vesicle size correlates linearly with the logarithm of RNA length.

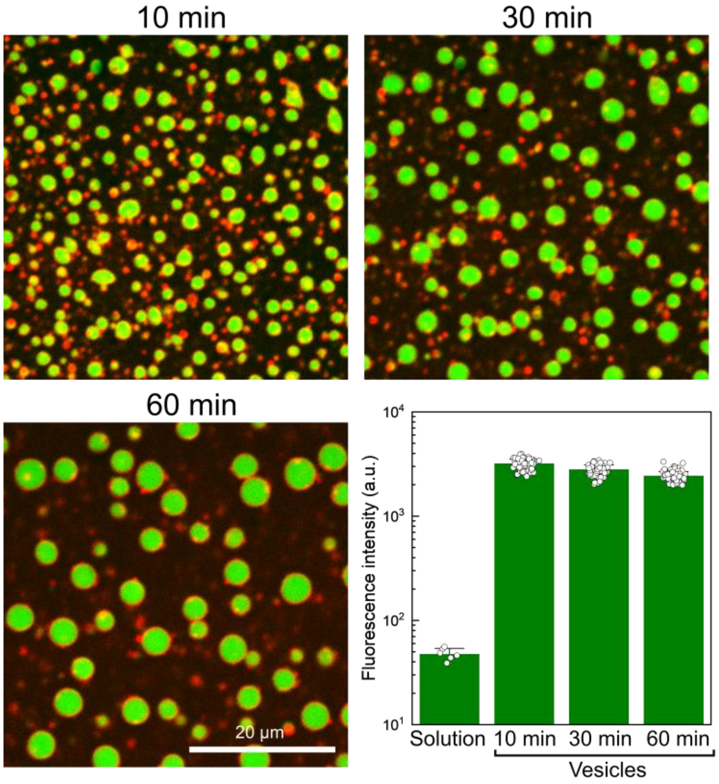

**Figure S10.** Confocal fluorescence microscopy images of 976 nt RNA-enriched vesicles after 10, 30, and 60 min. Quantifluor (green) and DiI (red) denote RNA and membrane signals, respectively. The graph shows average Quantifluor intensity in bulk solution and within vesicles (n = 100). RNA fluorescence inside vesicles is enriched 67 ± 6.7 fold relative to solution at 10 min and decreases gradually to 59 ± 6.0 fold after 30 min and 51 ± 4.8 fold after 60 min (76% of the 10 min value).

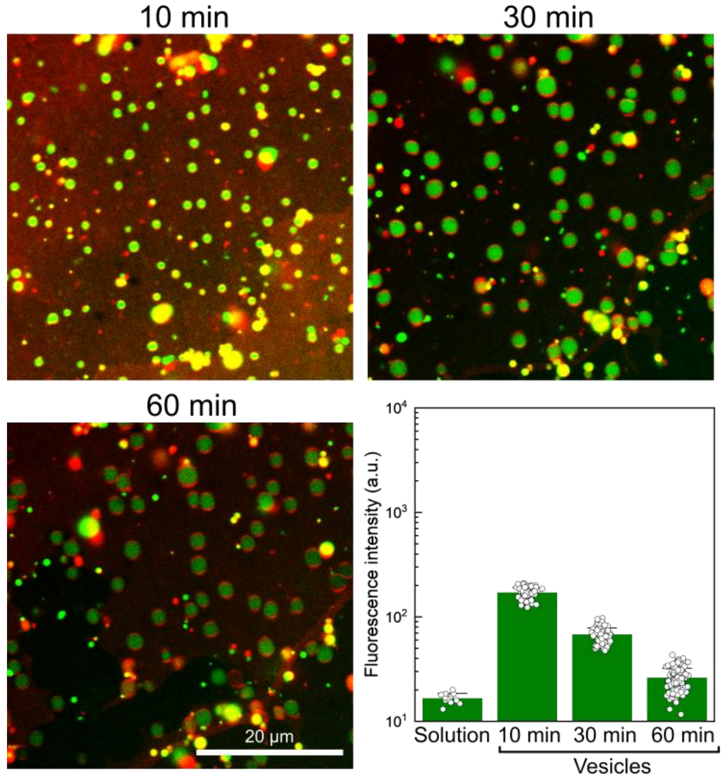

**Figure S11.** Confocal fluorescence microscopy images of 22 nt RNA-enriched vesicles after 10, 30, and 60 min. Quantifluor (green) and DiI (red) denote RNA and membrane signals, respectively. The graph shows average Quantifluor intensity in bulk solution and within vesicles (n = 100). RNA fluorescence inside vesicles is enriched 10 ± 1.1 fold relative to solution at 10 min and decreases gradually to 4.1 ± 0.6 fold after 30 min and 1.6 ± 0.4 fold after 60 min (15% of the 10 min value).

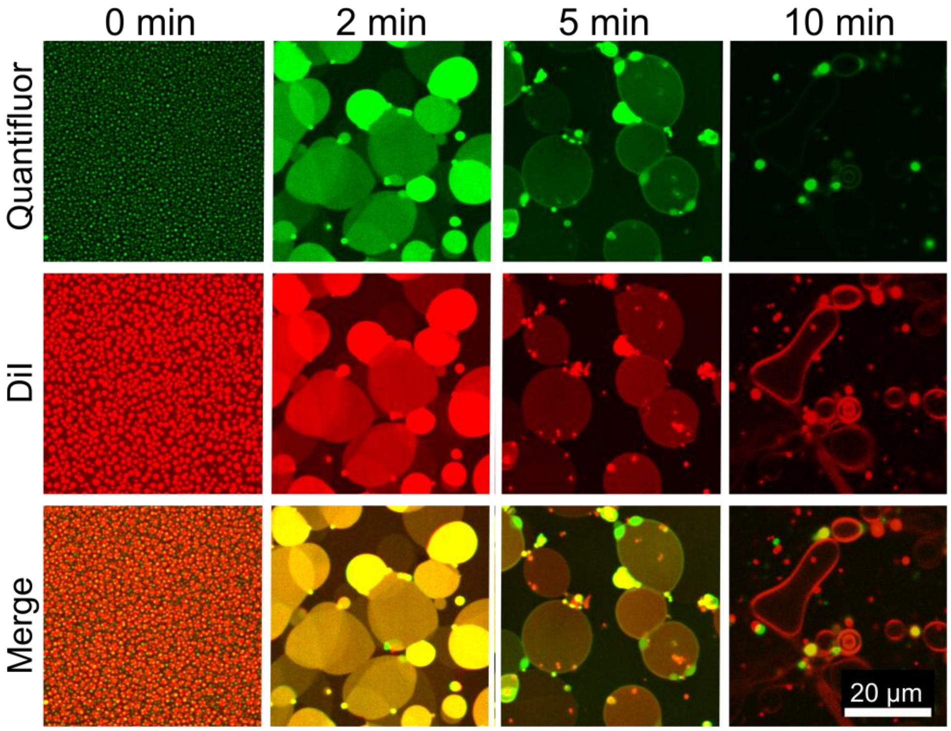

**Figure S12**. Time-lapse confocal fluorescence microscopy images of 10 nt RNA-$C_{12}$CT condensates following cysteine treatment.

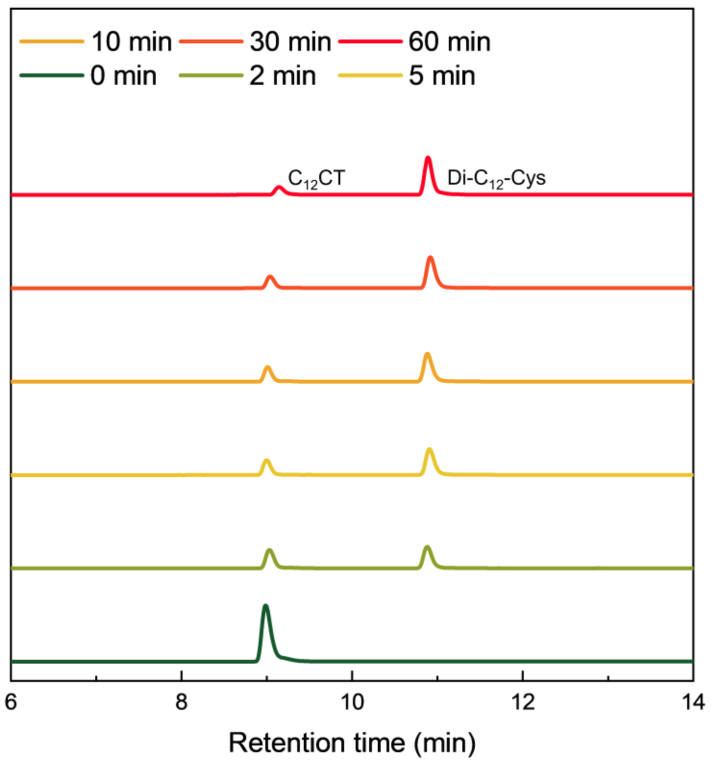

**Figure S13.** LCMS measurement of $C_{12}CT$ and diacylated cysteine (Di-$C_{12}$-Cys) after 10 nt RNA-$C_{12}CT$ condensates are treated with cysteine.

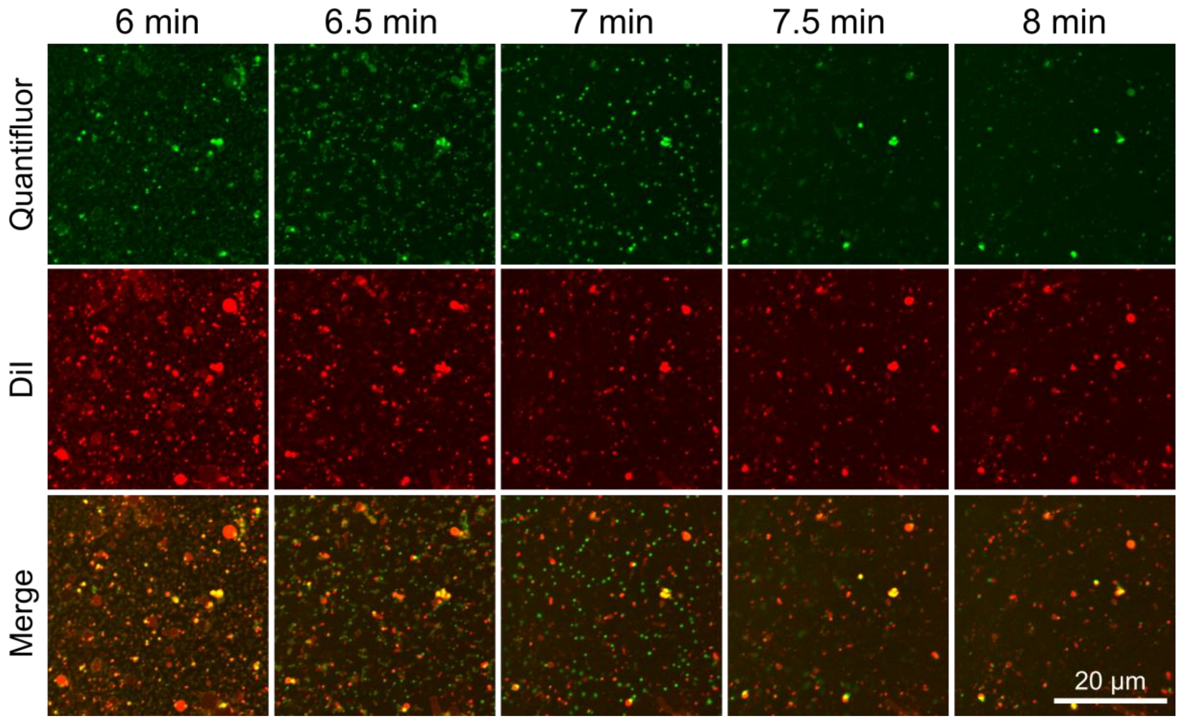

**Figure S14.** Time-lapse confocal fluorescence microscopy images of 15 nt RNA-$C_{12}$CT condensates following cysteine addition. Bright RNA-rich droplets appeared at 7 min but disappeared within 30 s, with no RNA-enriched vesicles observed thereafter.

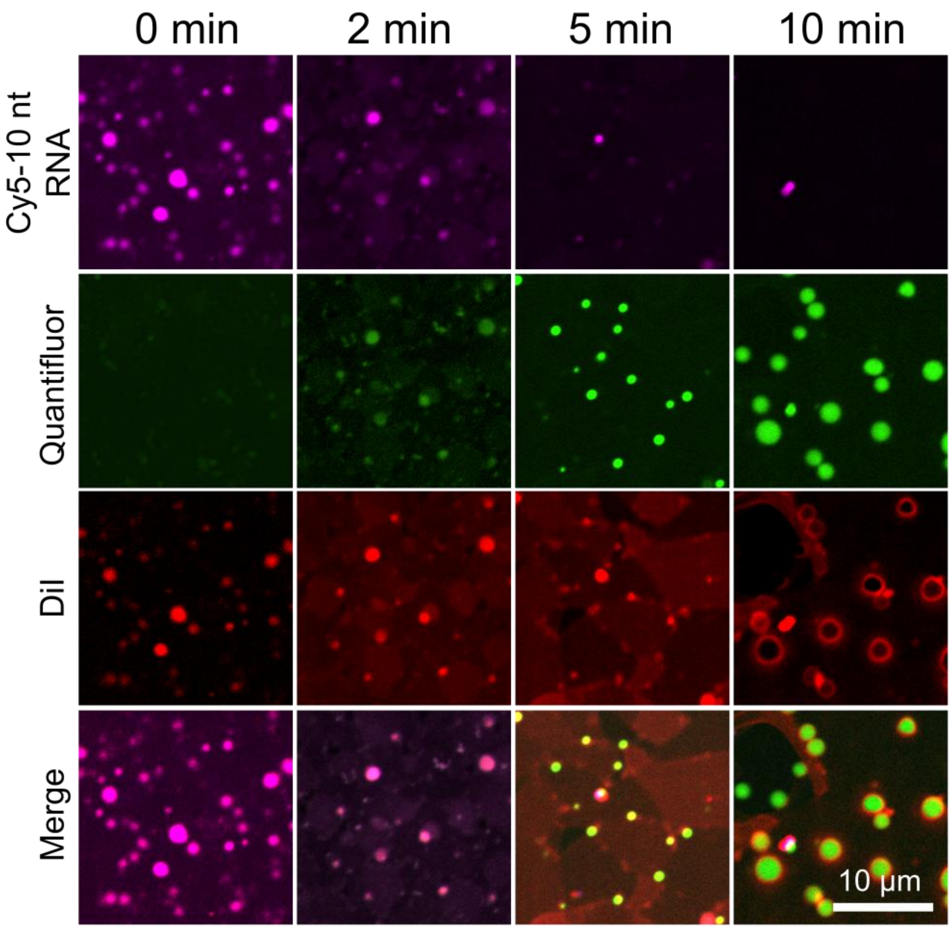

**Figure S15.** Time-lapse confocal fluorescence microscopy images of RNA-$C_{12}$CT condensates containing a 1:1 (w/w) mixture of Cy5-10 nt RNA and 80 nt RNA following cysteine addition.

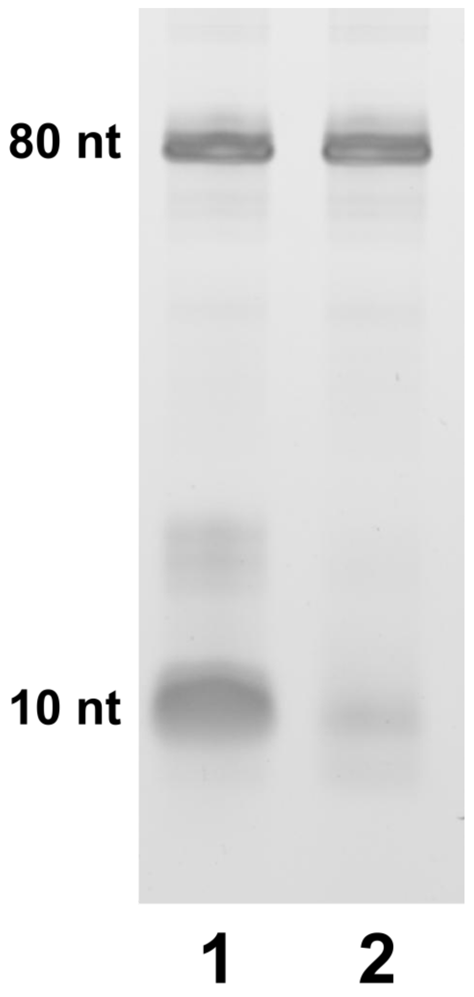

**Figure S16.** Urea-PAGE analysis of RNA-enriched vesicles formed from a 1:1 (w/w) mixture of 10 nt and 80 nt RNAs (1) in the absence and (2) in the presence of RNase A. Vesicles were treated with RNase A ($10^{-5}$ mg mL$^{-1}$) and incubated for 10 min at 37 °C.

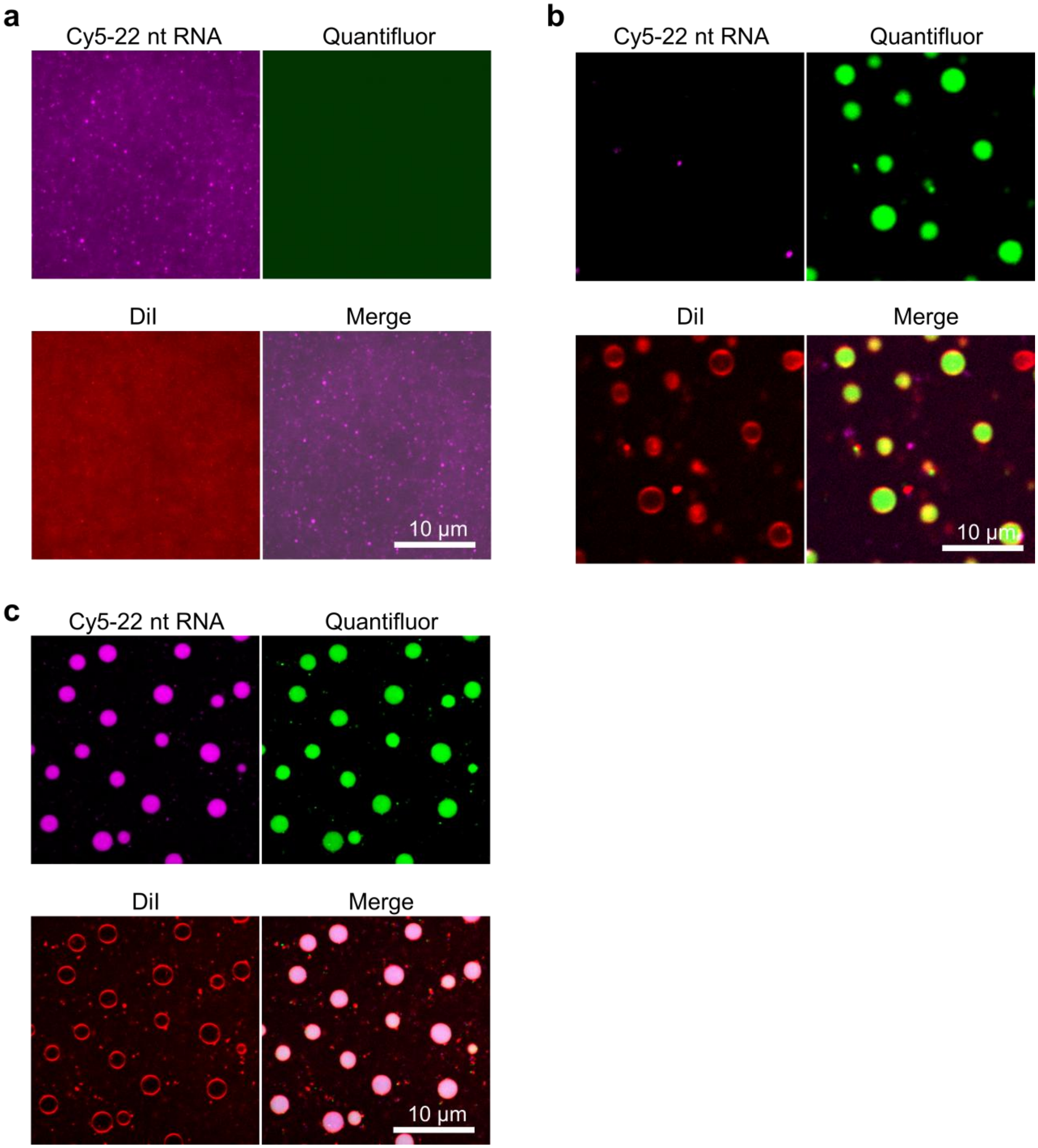

**Figure S17.** Confocal fluorescence microscopy images of RNA-$C_{12}$CT condensates containing a 1:1 (w/w) mixture of Cy5-22 nt RNA and 976 nt RNA, and of the corresponding RNA-enriched vesicles formed under different cysteine conditions. a) RNA-$C_{12}$CT condensates. b) RNA-enriched vesicles formed from the corresponding condensates 20 min after addition of 20% cysteine relative to the original condition. c) RNA-enriched vesicles formed from condensates of the same composition 10 min after addition of the original cysteine concentration (10 mM).

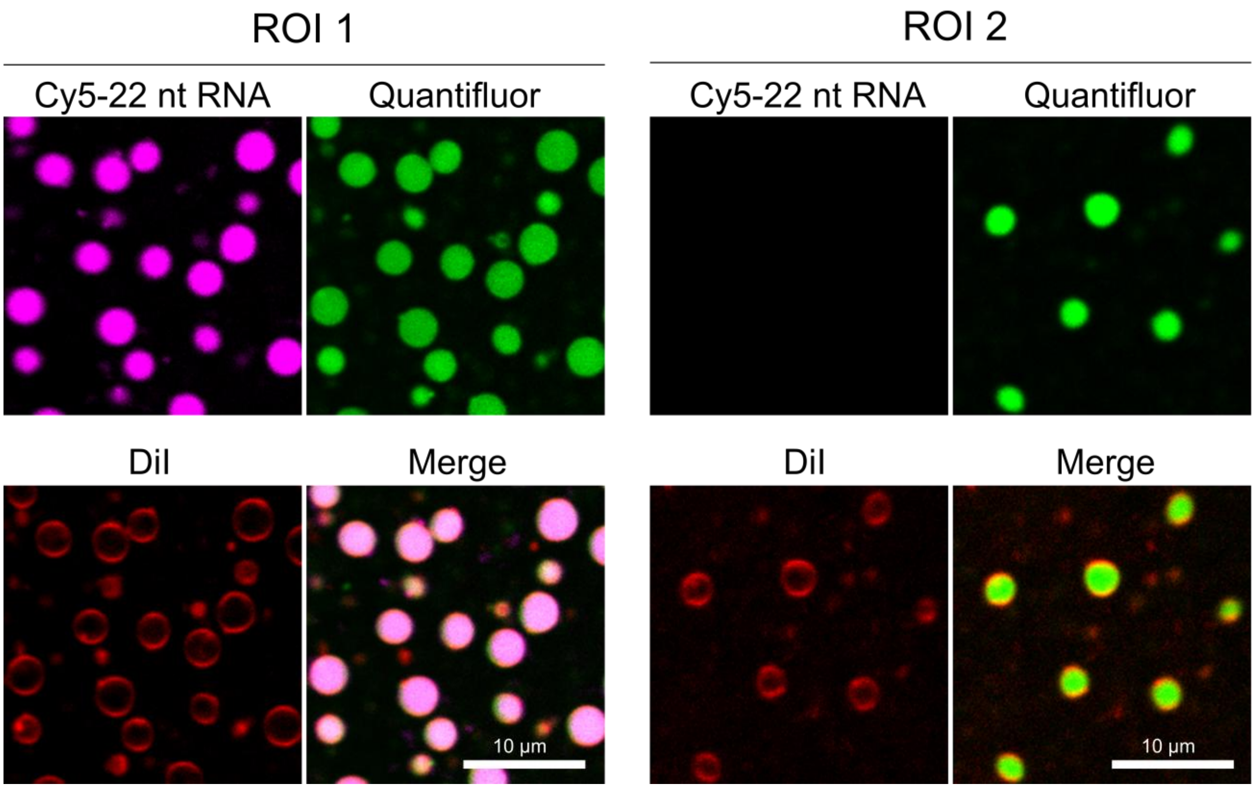

**Figure S18.** Individual fluorescence channel images of ROI 1 and ROI 2 shown in Fig. 3e.

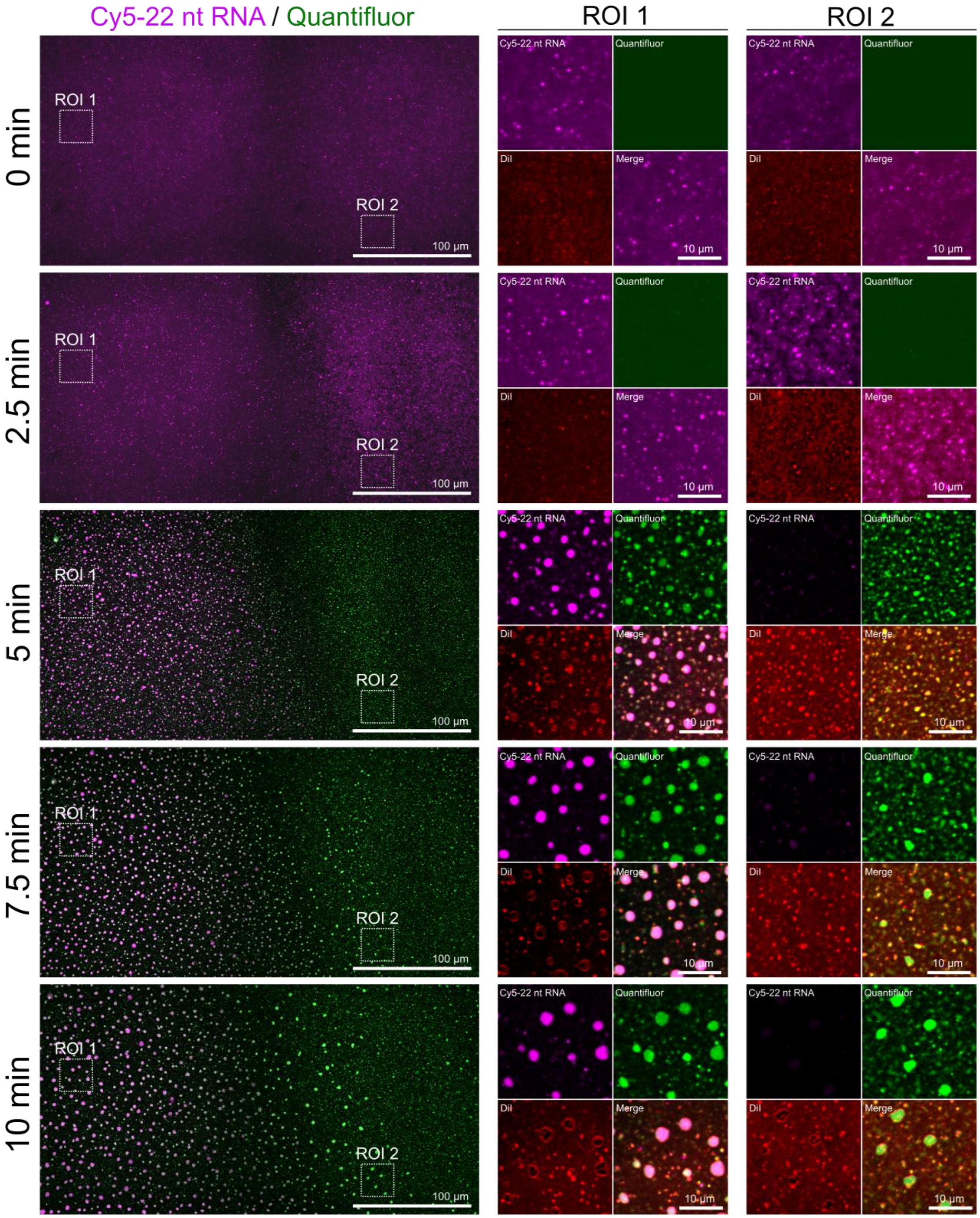

**Figure S19.** Time-lapse confocal fluorescence microscopy images of RNA-$C_{12}$CT condensates containing a 1:1 (w/w) mixture of Cy5-22 nt RNA and 976 nt RNA after addition of cysteine to the left edge of the chamber to establish a left-to-right cysteine gradient. ROI 1 and ROI 2 show magnified views near the left and right edges of the field of view, respectively, at each time point. At 0 min, Cy5-22 nt RNA and Quantifluor signals were homogeneously distributed, indicating uniform mixing of 22 nt and 976 nt RNAs. At 5 min, vesicles containing both Cy5-22 nt RNA and 976 nt RNA were observed near the left edge (ROI 1), whereas Cy5-22 nt RNA signal was lost near the right edge (ROI 2), where RNA condensates containing only 976 nt RNA were observed, as indicated by Quantifluor signal colocalized with DiI. At 7.5 min, vesicles enriched only in 976 nt RNA formed near the right edge (ROI 2), as indicated by strong Quantifluor signal in the absence of Cy5 signal and by DiI signal outlining the vesicle periphery. These time-lapse observations show that spatial patterning emerges during the condensate-to-vesicle transition owing to differences in vesicle-formation kinetics across the system.

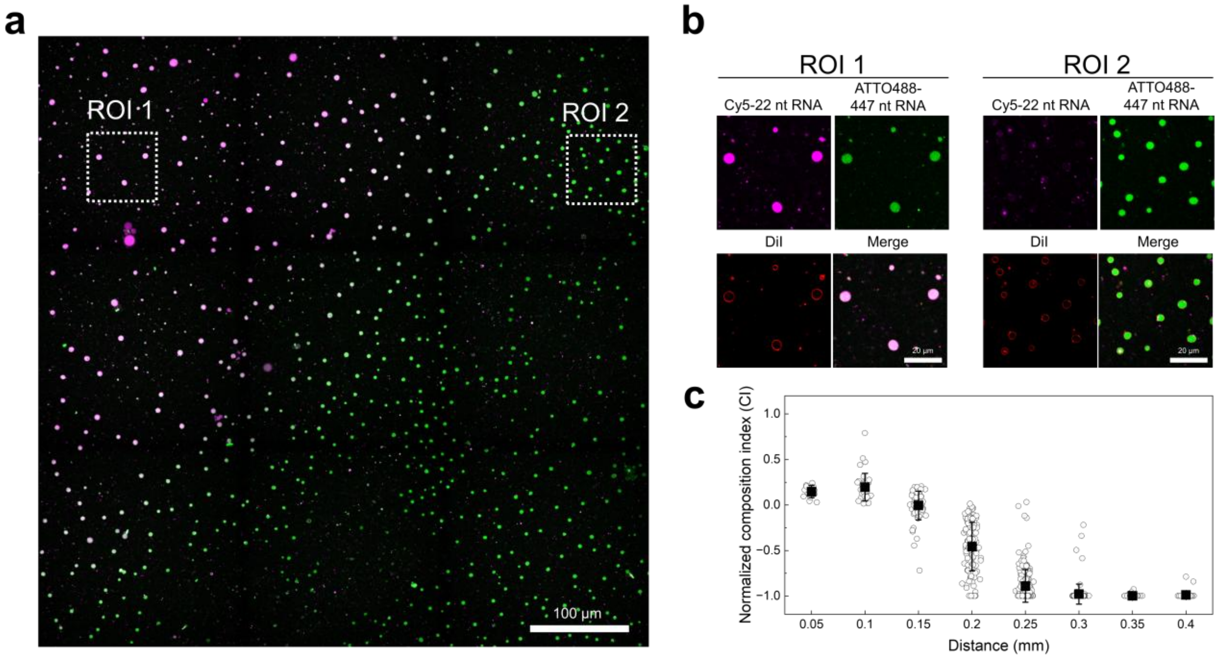

**Figure S20.** Spatial patterning of vesicles with distinct RNA compositions from a mixture of 22 nt and 447 nt RNAs under a cysteine gradient. a) Confocal fluorescence microscopy image of spatially patterned vesicles with distinct RNA compositions, acquired 20 min after cysteine addition to RNA-$C_{12}$CT condensates containing a 1:1 (w/w) mixture of Cy5-22 nt RNA and ATTO488-447 nt RNA. Cysteine solution was added to the left edge of the chamber to establish a left-to-right cysteine gradient. Magenta and green channels correspond to Cy5-22 nt RNA and ATTO488-447 nt RNA, respectively. The DiI channel is omitted for clarity. b) Magnified views of ROI 1 and ROI 2 in a, showing vesicles near the left and right edges of the image, respectively. c) Normalized composition index (CI) showing the relative intensities of Cy5-22 nt RNA and ATTO488-447 nt RNA fluorescence in a. CI values of 1 and −1 indicate the exclusive presence of Cy5-22 nt RNA signal and ATTO488-447 nt RNA signal, respectively. Distance on the x axis indicates distance from the left edge of the field of view shown in panel a.

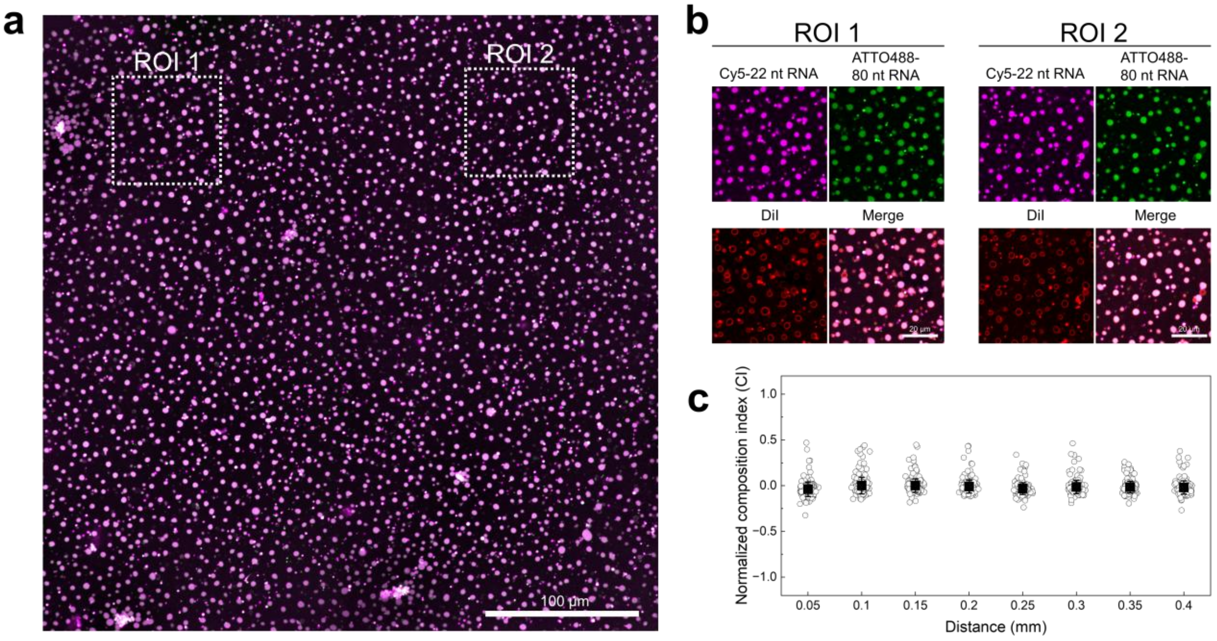

**Figure S21.** Spatially uniform vesicle formation from a mixture of 22 nt and 80 nt RNAs under a cysteine gradient. a) Confocal fluorescence microscopy image of vesicles with uniform RNA compositions, acquired 20 min after cysteine addition to RNA-$C_{12}$CT condensates containing a 1:1 (w/w) mixture of Cy5-22 nt RNA and ATTO488-80 nt RNA. Cysteine solution was added to the left edge of the chamber to establish a left-to-right cysteine gradient. Magenta and green channels correspond to Cy5-22 nt RNA and ATTO488-80 nt RNA, respectively. The DiI channel is omitted for clarity. b) Magnified views of ROI 1 and ROI 2 in a, showing vesicles near the left and right edges of the image, respectively. c) Normalized composition index (CI) showing the relative intensities of Cy5-22 nt RNA and ATTO488-80 nt RNA fluorescence in a. CI values of 1 and −1 indicate the exclusive presence of Cy5-22 nt RNA signal and ATTO488-80 nt RNA signal, respectively. Distance on the x axis indicates distance from the left edge of the field of view shown in panel a.

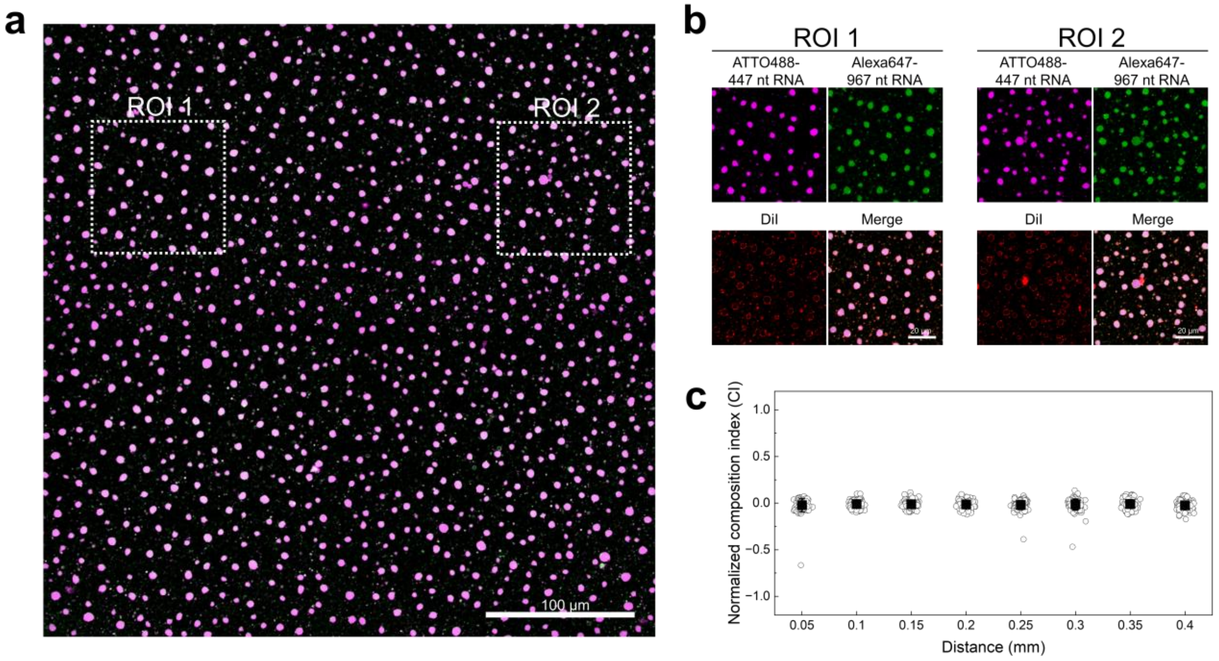

**Figure S22.** Spatially uniform vesicle formation from a mixture of 447 nt and 967 nt RNAs under a cysteine gradient. a) Confocal fluorescence microscopy image of vesicles with uniform RNA compositions, acquired 20 min after cysteine addition to RNA-$C_{12}$CT condensates containing a 1:1 (w/w) mixture of ATTO488-447 nt RNA and Alexa647-967 nt RNA. Cysteine solution was added to the left edge of the chamber to establish a left-to-right cysteine gradient. Magenta and green channels correspond to ATTO488-447 nt RNA and Alexa647-967 nt RNA, respectively. The DiI channel is omitted for clarity. b) Magnified views of ROI 1 and ROI 2 in a, showing vesicles near the left and right edges of the image, respectively. c) Normalized composition index (CI) showing the relative intensities of ATTO488-447 nt RNA and Alexa647-967 nt RNA fluorescence in a. CI values of 1 and −1 indicate the exclusive presence of ATTO488-447 nt RNA signal and Alexa647-967 nt RNA signal, respectively. Distance on the x axis indicates distance from the left edge of the field of view shown in panel a.

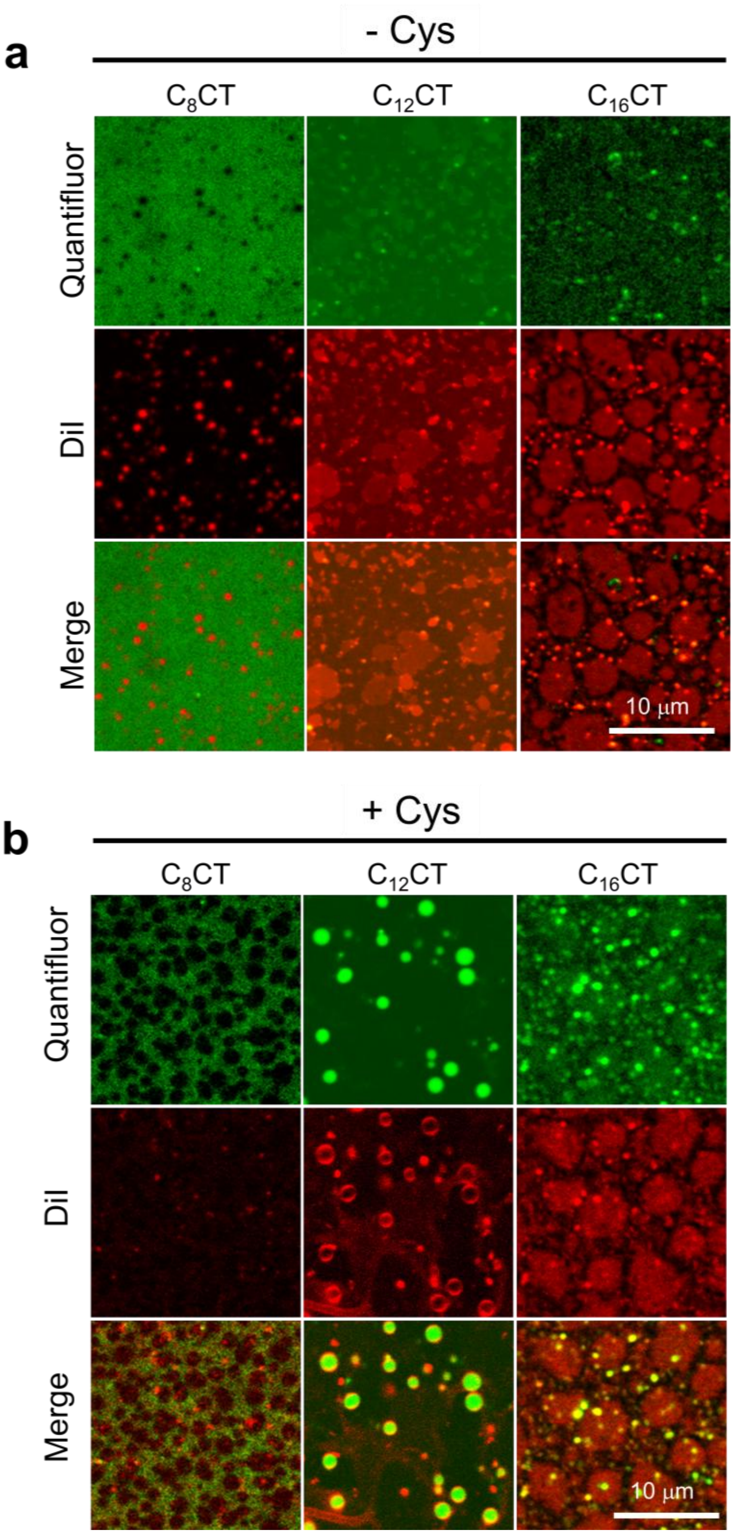

**Figure S23.** a) Confocal fluorescence images of 80 nt RNA-CT condensates formed using CT derivatives with varying aliphatic chain lengths: octyl ($C_8CT$), dodecyl ($C_{12}CT$), and hexadecyl ($C_{16}$CT) choline thioesters. b) Images obtained 10 min after cysteine treatment of the RNA-CT condensates shown in panel a. The experiments were conducted at 37 ˚C.

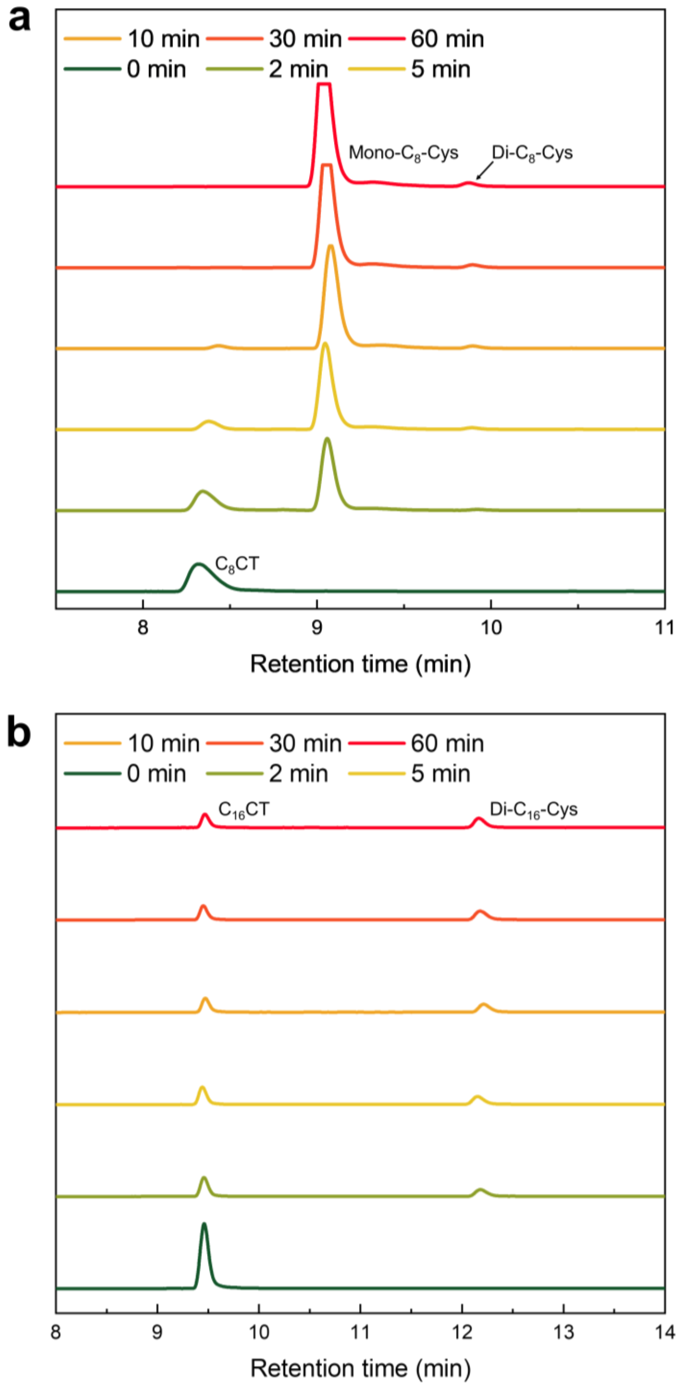

**Figure S24.** LCMS measurement of thioester and the corresponding mono- or diacylated lipids for a) 80 nt RNA-$C_8$CT condensates and b) 80 nt RNA-$C_{16}$CT condensates after addition of cysteine.

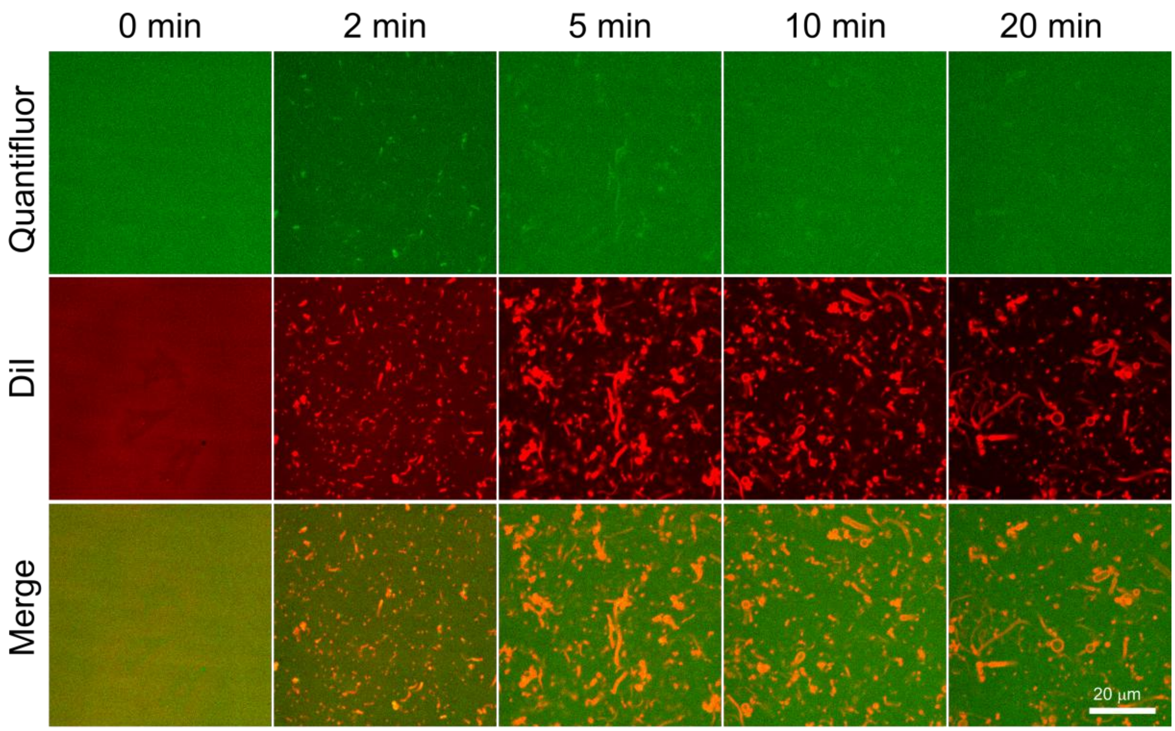

**Figure S25.** Time-lapsed confocal fluorescence microscopy images of 80 nt RNA-$C_8CT$ condensates after addition of 0.5 equivalent of cysteine relative to 4 mM $C_8CT$.

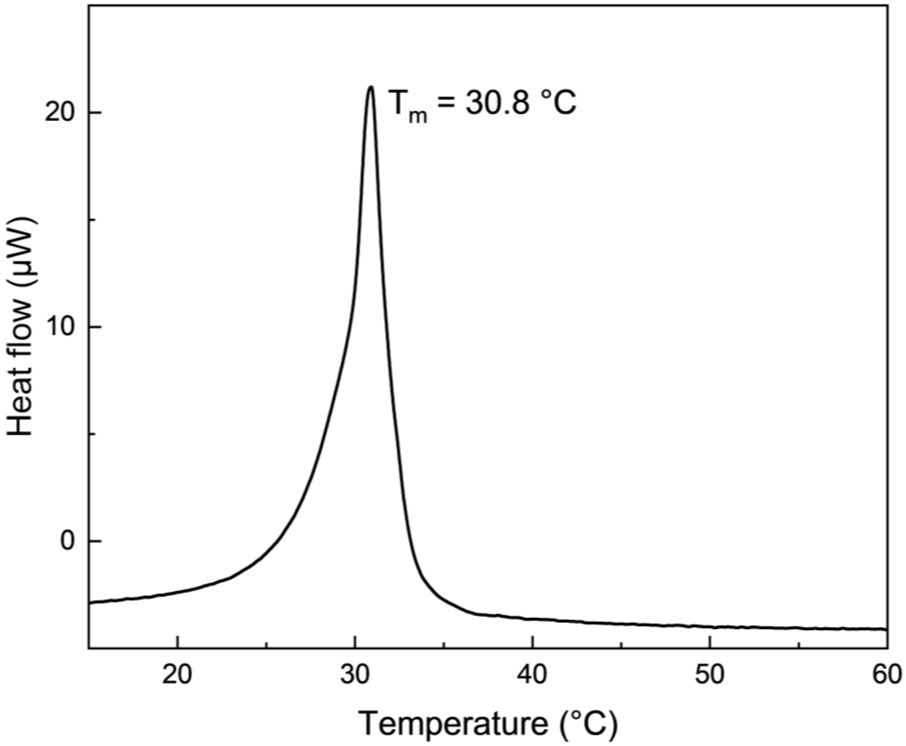

**Figure S26.** DSC thermogram of Di-$C_{12}$-Cys showing a phase transition at 30.8 °C.

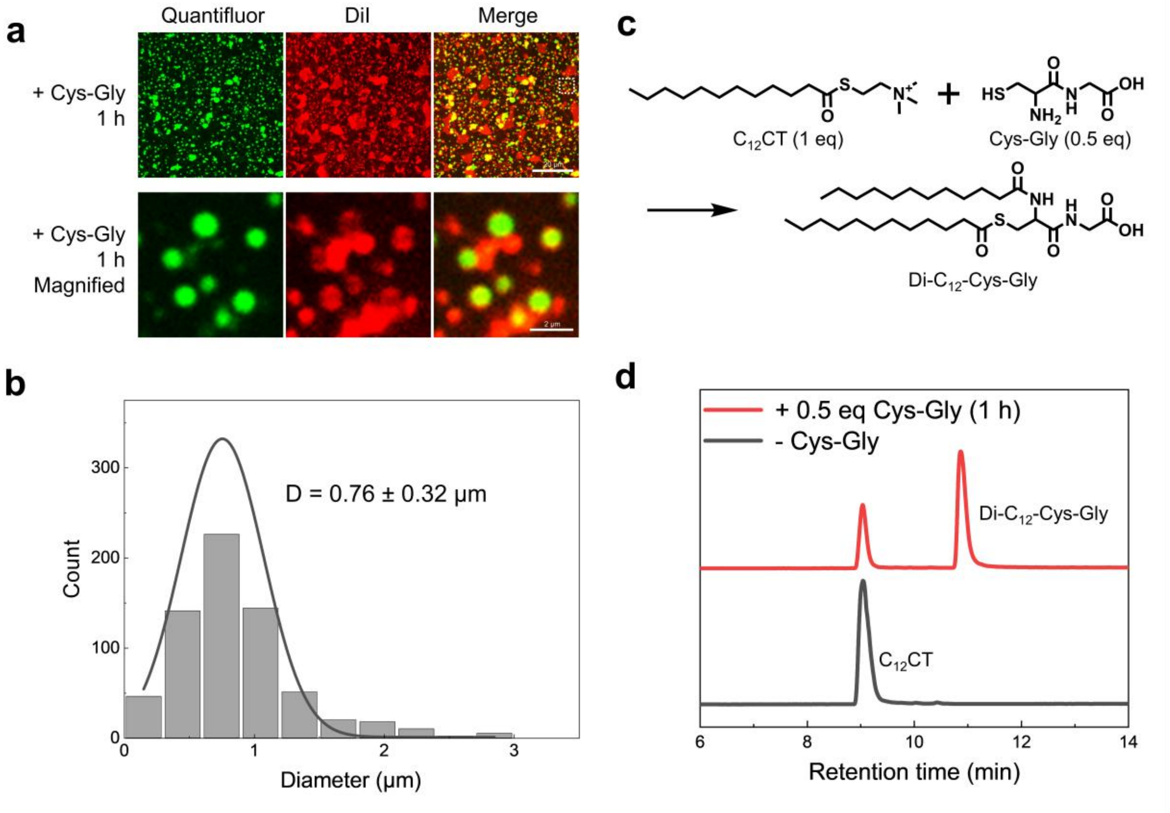

**Figure S27.** RNA-enriched vesicles generated using Cys-Gly. a) Confocal fluorescence microscopy images of RNA-enriched vesicles observed 1 h after addition of Cys-Gly to 80 nt RNA-$C_{12}$CT condensates. b) Size distribution of vesicles measured at 1 h, showing an average diameter of 0.76 ± 0.32 μm (n = 675). c) Schematic of the diacylation reaction between $C_{12}$CT and Cys-Gly producing Di-$C_{12}$-Cys–Gly. d) LCMS analysis of $C_{12}$CT consumption and formation of Di-$C_{12}$-Cys–Gly 1 h after treatment of 80 nt RNA-$C_{12}$CT condensates with Cys-Gly.

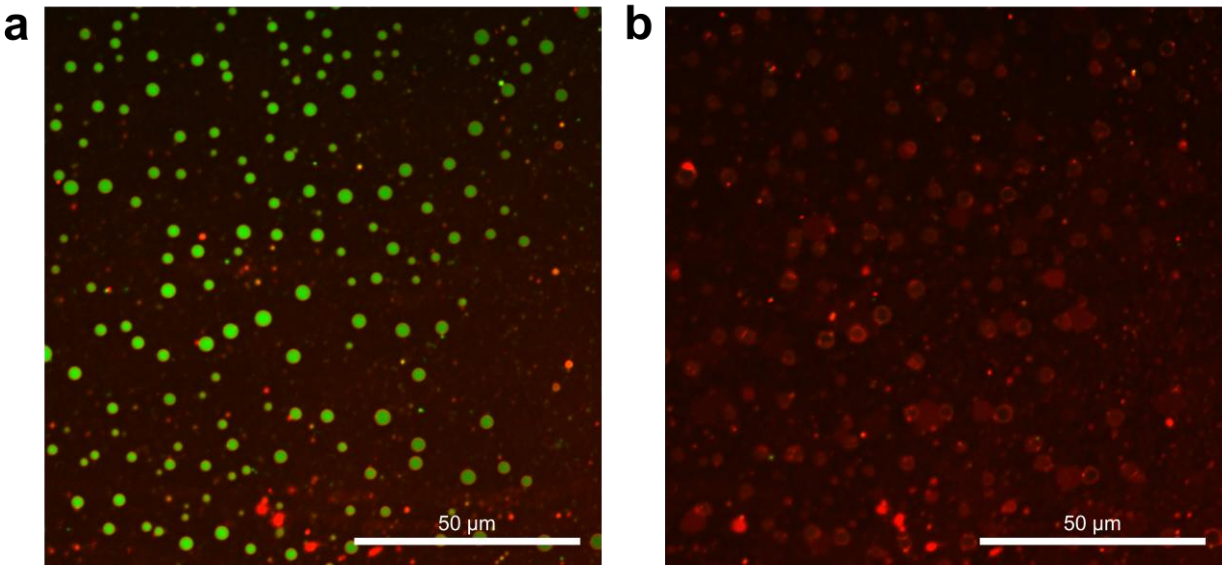

**Figure S28.** (a) Ribozyme and substrate RNA-enriched vesicles generated via a condensate-to-vesicle transformation. RNA solution (0.2 µM ribozyme and 3.3 µM substrate RNA in 120 µL of 20 mM HEPES, pH 8.2) was mixed with $C_{12}CT$ (100 µL, 11.8 mM in DIW) to form RNA-$C_{12}CT$ condensates, followed by treatment with cysteine (100 µL, 33 mM in DIW) to induce vesicle formation. The image was acquired 10 min after cysteine addition. (b) The same field of view as in (a) after 30 s exposure to 10 mM $MgCl_2$. Green and red signals correspond to Quantifluor RNA dye and DiI membrane dye, respectively.

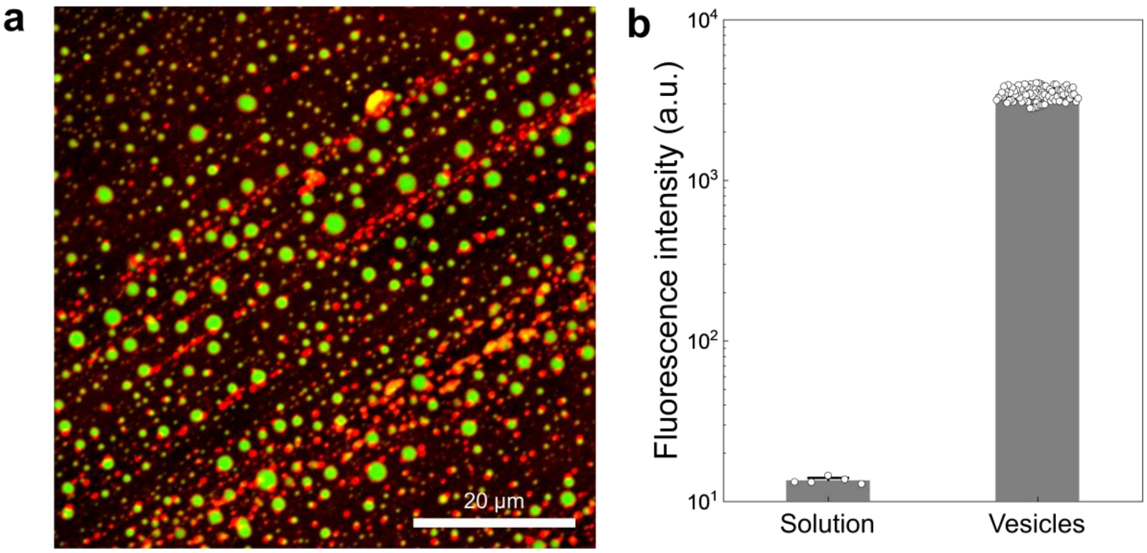

**Figure S29.** a) Representative confocal microscopic image of vesicles encapsulating ribozyme and substrate, acquired 20 min after cysteine addition to RNA-$C_{12}$CT condensates. Green signals indicate RNA stained with Quantifluor, and red signals indicate lipids stained with DiI. b) Quantifluor fluorescence intensity in bulk solution and within vesicles, quantified 10 min after cysteine addition to RNA-$C_{12}$CT condensates ($n = 85$). The vesicles exhibited a 260 fold higher fluorescence intensity than the bulk solution at 20 min incubation.

**a**

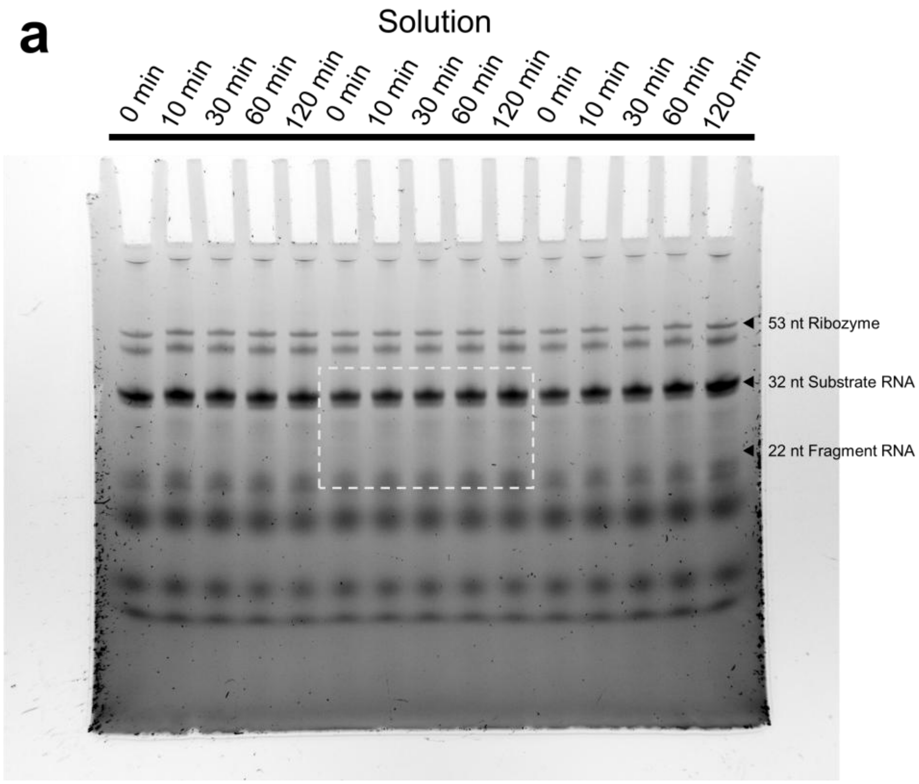
Solution
0 min
10 min
30 min
60 min
120 min
0 min
10 min
30 min
60 min
120 min
0 min
10 min
30 min
60 min
120 min
53 nt Ribozyme
32 nt Substrate RNA
22 nt Fragment RNA

**b**

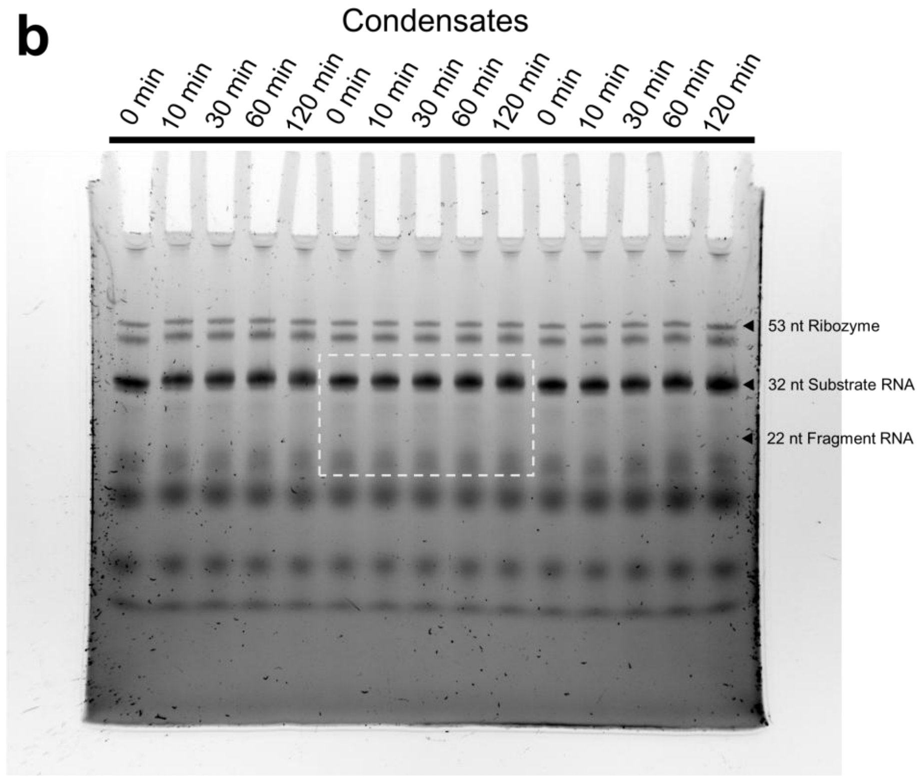
Condensates
0 min
10 min
30 min
60 min
120 min
0 min
10 min
30 min
60 min
120 min
0 min
10 min
30 min
60 min
120 min
53 nt Ribozyme
32 nt Substrate RNA
22 nt Fragment RNA

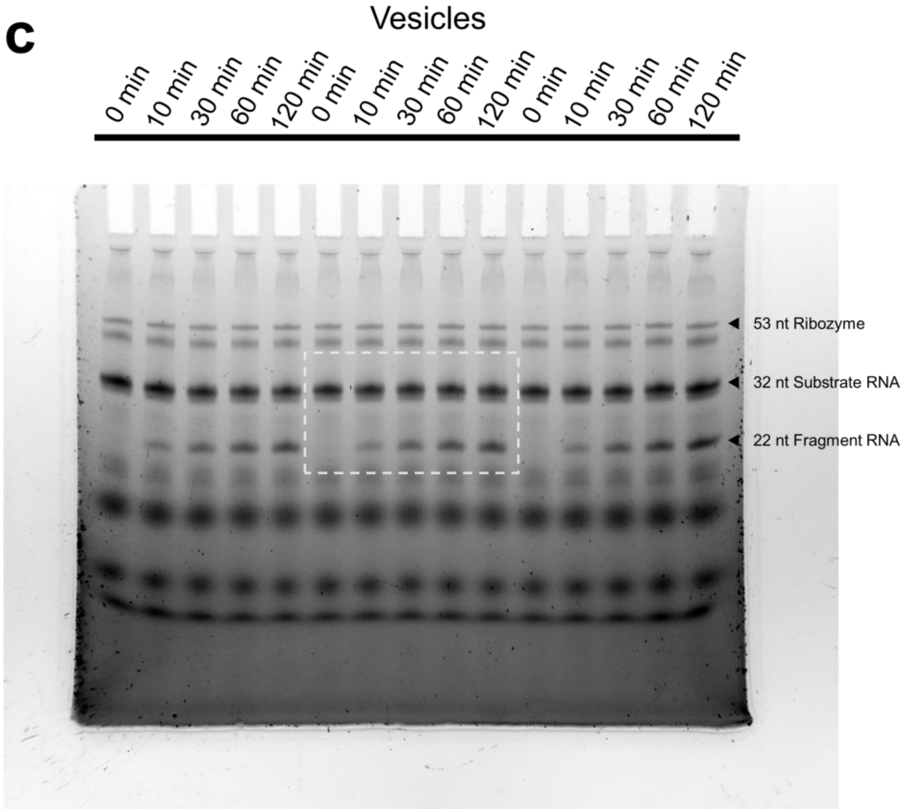

**Figure S30.** Uncropped gel images shown in Figure 4d. a) solution b) condensates c) vesicles. White dotted rectangle denotes the selection shown in the main text.

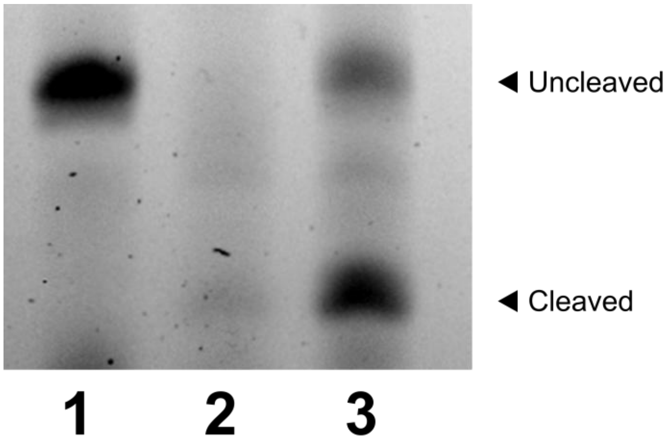

**Figure S31.** Urea–PAGE analysis of the hairpin ribozyme (R53) and substrate (S32) following 2 h incubation at 37 °C. 1) reaction mixture lacking $Mg^{2+}$, serving as a reference for the initial amount of S32. 2) reaction mixture containing $Mg^{2+}$ (0.1 mM) and RNase A ($10^{-5}$ mg $mL^{-1}$), showing extensive RNA hydrolysis. 3) RNA-enriched vesicles supplemented with $Mg^{2+}$ (0.1 mM) and RNase A ($10^{-5}$ mg $mL^{-1}$), showing the persistence of S32 and the appearance of the cleavage product F22, consistent with ribozyme activity inside the vesicles.

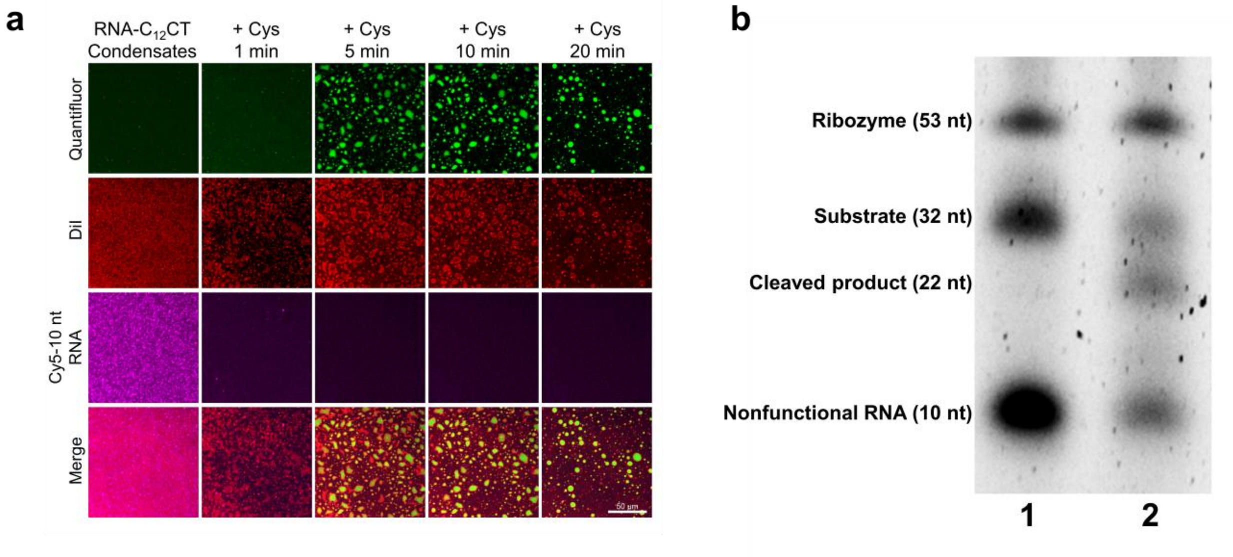

**Figure S32.** Selective partitioning of functional RNAs from mixtures of functional and nonfunctional RNAs. a) Time-lapse confocal fluorescence images showing selective enrichment and encapsulation of the 53 nt ribozyme and 32 nt substrate RNAs (Quantifluor signal), while excluding nonfunctional Cy5-10 nt RNAs during vesicle formation. RNA-$C_{12}$CT condensates were generated using final concentrations of 78 nM ribozyme, 1.3 μM substrate, 1.2 μM Cy5-10 nt RNA, and 0.1 mM $Mg^{2+}$. b) Urea-PAGE analysis of RNA-enriched vesicles formed from mixtures of functional and nonfunctional RNAs. Identical RNA compositions were used except that unlabeled 10 nt RNA replaced Cy5-10 nt RNA. (1) RNA mixture in solution without RNase A, representing the initial relative amounts of RNA species. (2) RNA-enriched vesicles treated with RNase A ($10^{-5}$ mg $mL^{-1}$). RNase A was added 20 min after cysteine treatment, followed by a 10 min incubation prior to analysis.

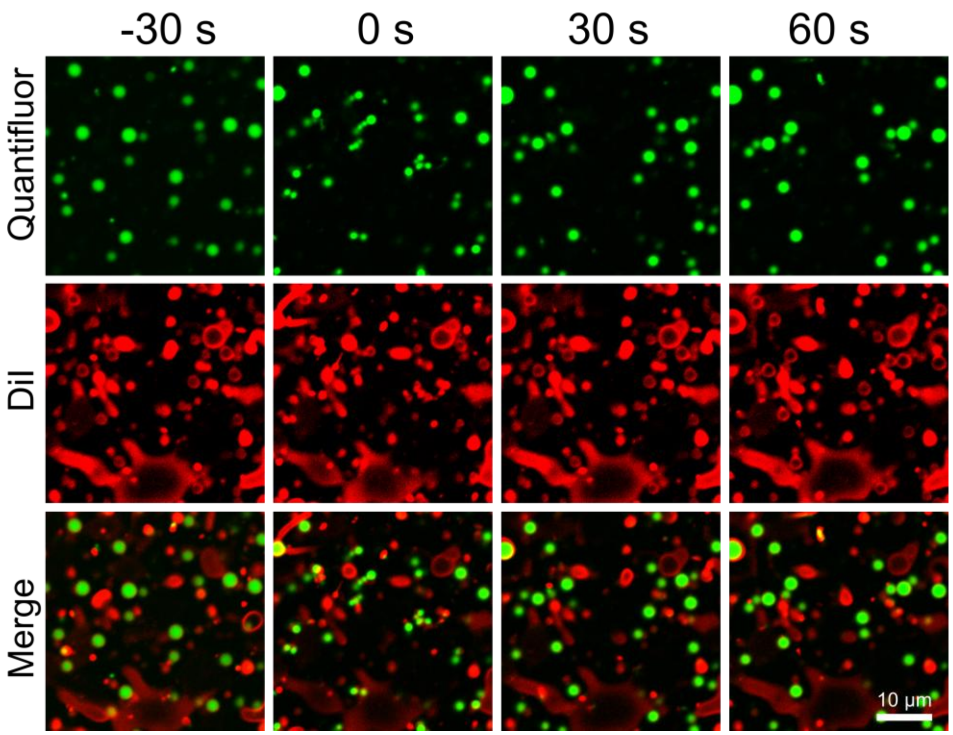

**Figure S33.** Time-lapse confocal fluorescence microscopy images of 80 nt RNA-enriched vesicles following addition of 4 M NaCl to achieve a final concentration of 400 mM (0 s). Vesicles initially shrank in response to increased ionic strength and transiently exhibited pearling (0 s), but gradually recovered their spherical morphology and original size within 60 s.

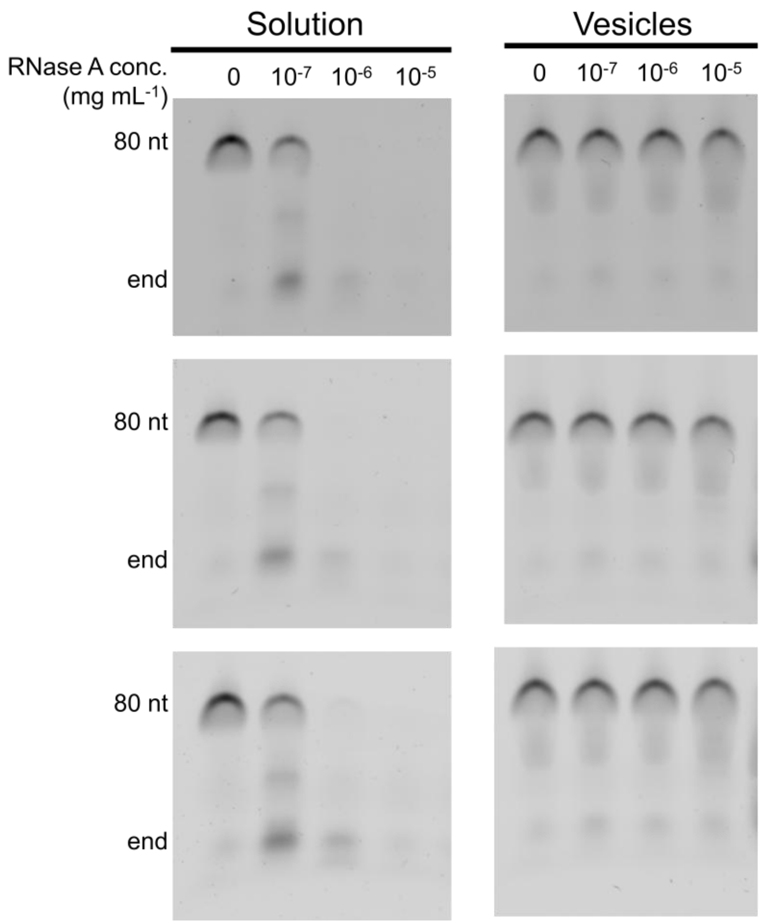

**Figure S34.** Urea–PAGE analysis of 80 nt RNA in bulk solution and in 80 nt RNA-enriched vesicles across a range of RNase A concentrations (0, $10^{-7}$, $10^{-6}$, and $10^{-5}$ mg $mL^{-1}$).
